# Synaptic Organization of Surface AMPARs Changes by Brain Region and Tauopathy

**DOI:** 10.1101/2024.07.22.604547

**Authors:** Rohit M. Vaidya, Jiahao Zhang, Duncan Nall, Rujuta Pendharkar, Yongjae Lee, Eung Chang Kim, Donghan Ma, Fang Huang, Hiroshi Nonaka, Shigeki Kiyonaka, Itaru Hamachi, Hee Jung Chung, Paul R. Selvin

## Abstract

The distribution of synaptic and extra-synaptic AMPA receptors (AMPARs) on neuronal plasma membranes is correlated with learning and memory. Although AMPAR organization has been extensively studied in neuronal cultures, its native cell-surface distribution in intact adult brain tissue across distinct brain regions and in neurodegenerative pathology remains poorly understood. Here, we combine a selective small-molecule labeling strategy with two-color 3D super-resolution dSTORM imaging to map native surface AMPAR organization at the nanoscale in 30 micron thick mouse brain slices. We find that wild-type mice exhibit marked regional differences in AMPAR organization, with the CA1 hippocampus containing a substantially larger extrasynaptic AMPAR pool than the nearby motor and somatosensory cortex. In the PS19 tauopathy mouse model, at an age preceding overt neurodegeneration, AMPAR organization is selectively disrupted in the hippocampus but largely preserved in the cortex. Specifically, we observe depletion of the extrasynaptic receptor pool together with reduced synaptic nanodomain organization, revealing early molecular-scale synaptic remodeling associated with tau pathology. These findings provide direct structural insight into region- and disease-dependent AMPAR organization in intact adult brain tissue and establish a broadly applicable framework for nanoscale investigation of synaptic receptor architecture in health and neurological disease.

## 1. Introduction

Synaptic plasticity plays a crucial role in learning and memory. ^1,2^ Among the diverse pre-and postsynaptic mechanisms that regulate synaptic function activity-dependent changes in the number, constitution, and conductivity of α-amino-3-hydroxy-5-methyl-4-isoxazolepropionic acid receptors (AMPARs) in the post-synaptic membrane are major contributors to excitatory synaptic plasticity in the mammalian brain.^3–5^ Thus, alteration in surface- (or cell membrane-) bound AMPARs at excitatory synapses can provide important molecular readouts of postsynaptic organization and synaptic strength. Furthermore, dysregulation of AMPAR signaling and trafficking has been implicated in multiple neurological diseases.^6–8^

The highly crowded nature of synaptic proteins and the small distance (∼20-30 nm) of the synaptic cleft at a chemical synapse ^9^ requires the use of high-resolution imaging techniques to study the intricate nanoscale organization of AMPARs. A variety of super-resolution fluorescence imaging techniques have been used to show that AMPARs are organized in post-synaptic nanodomains that are correlated with glutamate release sites at the pre-synaptic membrane, ^10–12^ forming a part of a trans-synaptic nano-column. ^13^ AMPARs are also present at extra-synaptic sites as a reserve pool and can transition between synaptic and extra-synaptic sites rapidly through lateral diffusion along with endocytic or exocytic trafficking during synaptic plasticity to regulate their number at the synapse in response to synaptic activity^.4,5,14,15^

Most of these studies, however, have been performed on primary neuronal cultures. While being a good model *in vitro* system, primary cultures lack the physiological 3D-brain circuitry critical for learning and memory which is affected by numerous neurologic diseases. ^16,17^ However, visualizing and discerning the AMPAR distribution in thick brain tissue is difficult. The first challenge is selective labeling of native surface AMPARs. Getting access to native surface AMPARs in neurons deep into the tissue is difficult with conventional antibodies. ^18^ To facilitate the labeling of surface AMPARs, previous studies have attached fluorescent proteins or tags on AMPAR, but these protein modification strategies can suffer from over-expression artifacts.^14,19–21^ Even in knock-in (KI) mice, ^22^ the presence of a bulky tag can hamper normal AMPAR function and trafficking. ^23^ A promising new approach uses KI mice with a small biotin acceptor peptide (AP) tag on the GluA2 subunit of AMPARs, but this is limited to GluA2 and requires an additional biotinylation step, making the labeling process complicated. ^24^ Another challenge is achieving high-resolution imaging of surface AMPARs in thick tissue. Depth-dependent optical aberrations and increased scattering due to refractive-index mismatch results in poor resolution. ^25,26^ Recent advances using adaptive optics and in-situ PSF retrieval (INSPR) methods have achieved <10 nm lateral and <30 nm axial localization precision, but their application has been limited to imaging a single color, and hence getting the distances between molecules has not been possible.^27–29^

In this work, we use a recently developed probe, called Chemical AMPAR Modification 2 (CAM2) bound to a fluorophore (Alexa Fluor 647) to label native surface AMPARs in mouse brain slices which are kept alive during labeling,^18,30,31^ CAM2 labels all four GluA1-4 subunits^30^. The small size of CAM2 allows the probe to diffuse rapidly throughout the 200 mm brain slice compared to conventional antibodies.^18^ Combining this labeling scheme with two-color 3D direct Stochastic Optical Reconstruction Microscopy (dSTORM) ^32,33^ that utilizes adaptive optics and INSPR, we observe the nanoscale organization of native AMPARs that are on the cell-surface as a function of brain regions and tauopathy, for the first time in mouse brain slices.

To observe brain region-specific distributions, we compared the CA1 region of the hippocampus (referred to as “hippocampus”) with the motor and somatosensory cortical regions (referred to as “cortex”) that lie directly above the hippocampus in coronal sections of 6-month-old Thy1-YFP-H mice brains (called WT). Despite the neighboring proximity of these two regions, differences in AMPAR nanoscale organization are detected, possibly indicating a molecular basis for their physiological differences. ^34,35^ To observe pathological changes, we examined 6-months old PS19 mouse model of Alzheimer’s disease exhibiting tauopathy, where we find disruptions in the distribution of AMPARs on both extra-synaptic and nanoscale synaptic levels. These results suggest a direct molecular-level basis for the learning deficits in these mice which are evident at this age, before the onset of neurodegeneration. ^36^

## 2. Results

### 2.1. Visualizing synaptic nanoscale architecture of native surface AMPARs in mouse brain slices

CAM2 utilizes ligand-directed acyl imidazole (LDAI) chemistry for covalent attachment of a small probe to all AMPAR subunits GluA1-4. 6-pyrollyl-7-trifluoromethyl-quinoxaline-2,3-dione (PFQX) is used as a ligand, which recognizes surface AMPARs and facilitates an acyl-substitution reaction of the labeling fluorophore to nucleophilic amino acid residues such as Lys, Ser or Tyr located near the ligand-binding domain. The fluorophore is thus directly attached to the subunit while the ligand moiety gets cleaved and can be washed away. CAM2 conjugated to Alexa Fluor 647 was used to fluorescently label surface AMPARs in 200 µm thick *live* acute brain slices from Thy1-YFP-H mice at 6 months of age (**Figure 1a**). Using this mouse line allows us to identify a sparse population of single pyramidal neurons and CAM2-labeled AMPARs associated with them. Keeping the brain slices alive during the labeling step ensures that only the surface AMPARs are being labeled (**Figure S1**). After the incubation with CAM2, acute brain slices were fixed and further cryo-sectioned down to 30 µm thickness before detergent-mediated permeabilization and immunostaining with CF568-labeled nanobody against PSD-95, or with primary and secondary antibodies against Homer-1. PSD-95 and Homer-1 are located intracellularly in the post-synaptic density (PSD) and help identify the location of excitatory synapses.

**Figure 1.**
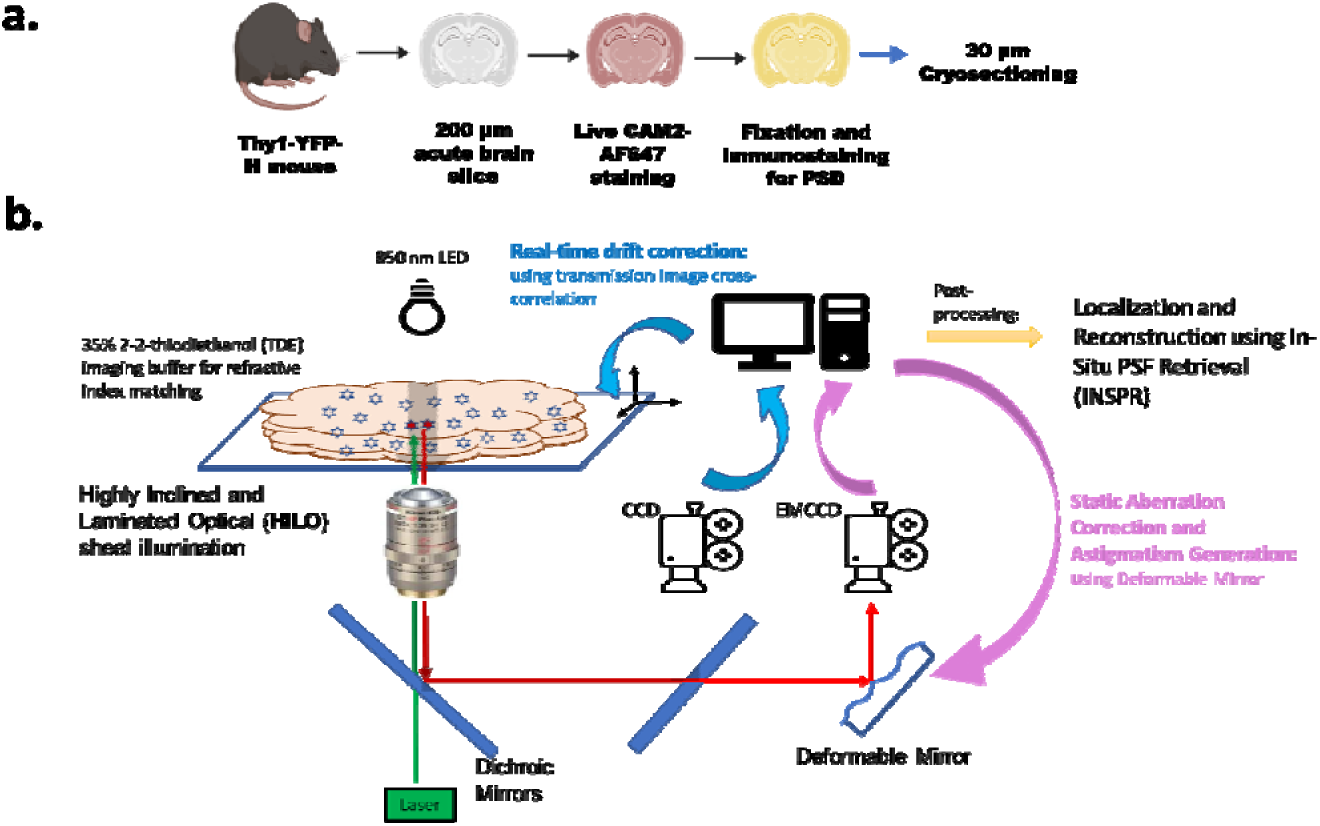
Labeling and Imaging Scheme. a, Schematic of the sample preparation 59. 200 µm acute brain slices fromThy1-YFPH mice are live-stained with CAM2-Alexa Fluor 647 followed by fixation and immunostaining for PSD and cryosectioning (Materials and Methods) b. Schematic of the microscope setup. Imaging in refractive index matched medium allowed reducing light scattering. Additional aberrations were corrected using a deformable mirror and in-situ PSF retrieval (INSPR). Real-time stage drift correction allowed for coarse drift correction, which was corrected further using cross-correlation during image processing.

For 3D dSTORM imaging of 30 µm cryosections with high resolution, we used a combination of strategies: refractive index matching for reducing light scattering, static aberration correction using a deformable mirror, and INSPR, and a 3-step drift correction involving real-time stage drift correction for coarse drift, followed by intra-channel and inter-channel 2D cross-correlation for finer drift.(**Figure 1b**). The deformable mirror was also used for generating astigmatism for axial information. This imaging scheme allowed us to achieve <10 nm lateral and <30 nm axial localization precisions throughout the 30 µm thick cryosections (**Figure S2 and S3**). For accurate two-color imaging, we measured and corrected chromatic aberrations as a function of imaging depth (Figure S2f).

Using this labeling and imaging scheme, we visualized native surface AMPARs and PSD-95 in Thy1-YFP-H mice brain slices. In the diffraction-limited images in the cortex, we observe surface AMPAR and PSD-95 colocalized with YFP spines (**Figure 2a**). With super-resolution 3D dSTORM imaging (Figure 2b), we observe single molecule blinking events (referred to as localizations) of surface AMPARs and PSD-95; their distributions were clearly resolved **(Figure 2c, d**). At 15 µm deep, the lateral localization precision was 7 nm, and the axial localization precision was 22 nm: a 3D view can be seen in the supplementary movie (**Figure SM1**). We used a modified DBSCAN algorithm to identify clusters of PSD-95 and clusters of AMPAR (**Figure 2e**), based on their localization density and minimum size ^37^. The average distance between AMPAR clusters and PSD-95 clusters was found to be 14 nm (**Figure 2k**, **Table S1**). We also separately labeled the PSD protein Homer-1 (**Figure 2g-j**). At 18 µm deep, the lateral resolution was 6 nm, and the axial resolution was 20 nm: a 3D view can be seen (**Figure SM2**). The distance between AMPAR clusters from Homer-1 clusters was found to be 64 nm (**Figure 2l**, **Table S2**). These values agree closely with previously reported distances of PSD-95 and Homer-1 from the post-synaptic membrane measured using electron microscopy ^38,39^ and dSTORM ^40^, thereby validating our labeling and imaging technique

**Figure 2.**
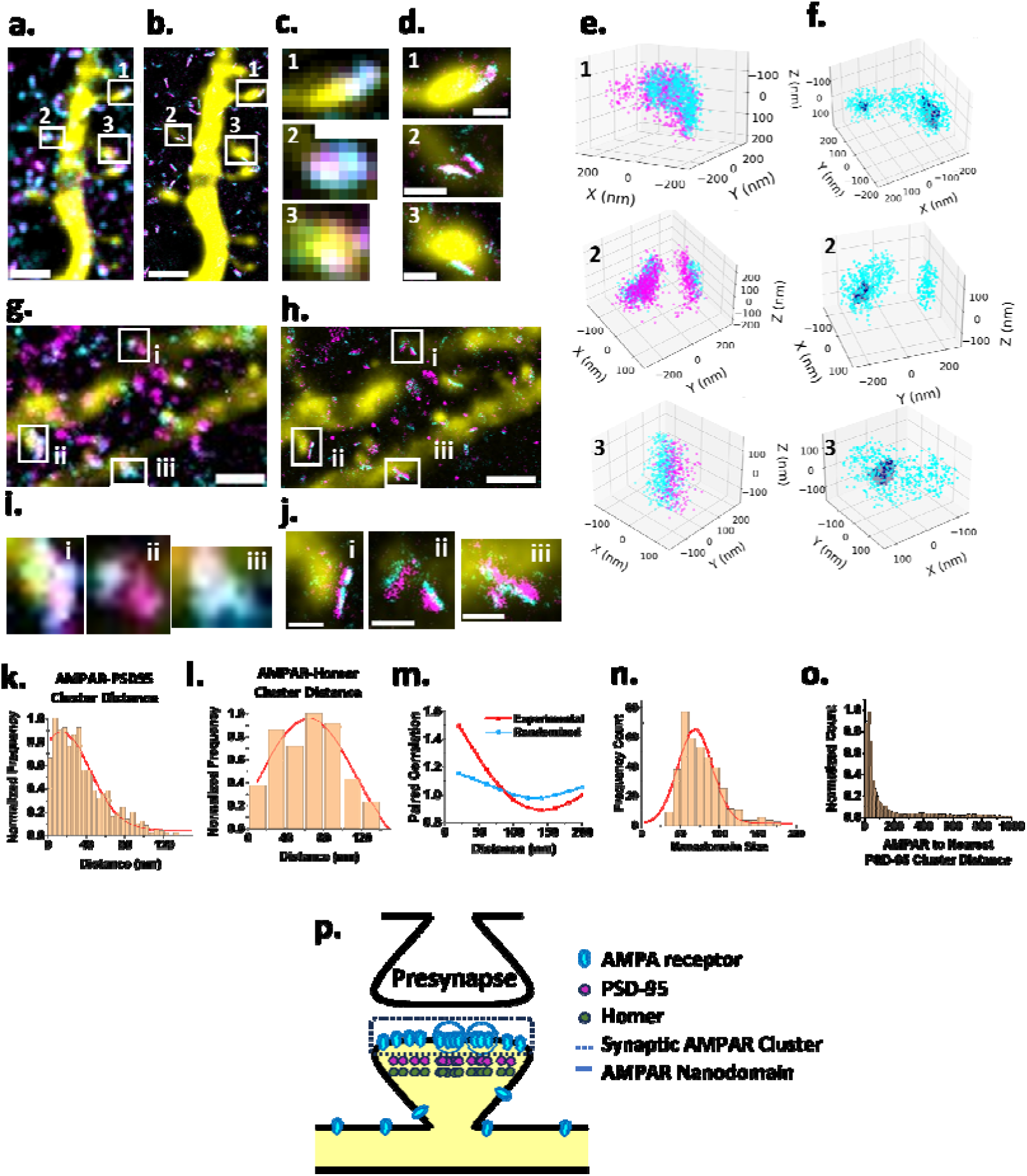
Nanoscale Organization of native surface AMPARs in mouse brain slices. **a**, Diffraction limited image of a YFP neuron (yellow); AMPAR (cyan) labeled with CAM2-Alexa647 and PSD-95 (magenta) labeled with a nanobody conjugated to CF568. **b,** Super-resolution dSTORM reconstruction of AMPAR and PSD-95 from b. **c,** Zoomed in ROIs on individual YFP spines (white boxes) in a. **d,** Zoomed in ROIs on individual YFP spines (white boxes) in b. **e,** AMPAR and PSD-95 clusters from the zoomed in ROIs in d. **f,** AMPAR nanodomains (navy blue) within the clusters in e. **g,** Diffraction limited image of a YFP neuron (yellow); AMPAR (cyan) labeled with CAM2-Alexa647 and Homer-1 (magenta). **h.** Super-resolution dSTORM reconstruction of AMPAR and PSD-95 from g. **i,** Zoomed in ROIs of individual YFP spines (white boxes) from g. **j.** Zoomed in ROIs of individual YFP spines (white boxes) from h. **k,** Distance between AMPAR-PSD-95 clusters (n=499 cluster pairs from the cortex from three 6-month-old Thy1-YFP-H mice). The peak of the fitted Gaussian function is at 14 nm with standard deviation of 30 nm (R-square=0.95). **l,** Distance between AMPAR-Homer-1 clusters (n=158 cluster pairs from one 6-month-old Thy1-YFP-H mouse. The peak of the fitted Gaussian function is at 64 nm with a standard deviation of 51 nm (R-square=0.90). **m,** Paired correlation function of AMPAR and PSD-95 localizations within a cluster pair (n= 499 pairs). Red: experimental data, Blue: simulated randomized distribution. **n,** AMPAR nanodomain size. The peak of the fitted Gaussian function is at 69 nm with a standard deviation of 22 nm (R-square=0.90). **o,** Distance of AMPAR localization from the nearest PSD-95 cluster (n=15 YFP cortical neurons from three 6-month-old Thy1-YFP-H mice). p, Schematic of the post-synaptic organization of AMPA receptor, PSD-95 and Homer.

The distribution of localizations within an AMPAR-cluster correlates with the corresponding PSD-95 cluster, as shown by comparison against simulated randomized distributions (**Figure 2m**, **Table S3**). Looking within the individual AMPAR clusters (**Figure 2e**), we observed high-density peaks of AMPAR, which were isolated using another modified DBSCAN algorithm (**Figure 2f**). Previous studies in dissociated neuronal cultures called these peaks “nanodomains” that form a part of the trans-synaptic nanocolumn with the pre-synaptic neurotransmitter release sites. ^10,13^ We found the size of AMPAR nanodomains to be 69 +/−22 nm (standard deviation) in our brain slices (**Figure 2n**, **Table S4**), matching closely with the previously reported values (∼ 69.6 nm, interquartile range of 54-93 nm)^10^ in dissociated neuronal cultures.

We next measured the distance of each localization of an AMPAR around a PSD-95 cluster along a YFP-positive cortical neuron, with the goal of determining synaptic vs extra-synaptic AMPARs (**Figure 2o**). For accurate AMPAR-to-PSD distance measurement, we measured the distance of each AMPAR localization within 1 µm of a PSD-95 cluster to the concave hull created around the PSD-95 cluster, The clustering parameters for PSD-95 were optimized to avoid false PSD-95 cluster detections due to any low density signals from the background and inaccurate PSD shape determination when generating the concave hulls (Methods). It should be noted that we detected PSD-95 clusters on about a third of total spines, thus there could be a potential bias in this analysis towards counting only the synapses with a sufficiently high degree of PSD-95 enrichment to be considered as PSD95 clusters. In order to ensure that the PSD-95 labeling is not affected by the CAM2-labeling of AMPARs, we compared PSD-95 labeling with and without CAM2 and labeled all AMPARs post-permeabilization with a conventional anti-GluA2/3 antibody instead. We did not find any effect on PSD-95 labeling due to CAM2 (**Figure S4 and S5**).

The peak of the AMPAR-to-PSD distance distribution lies at 20 nm, which corresponds with the previously measured distance between PSD-95 and AMPAR clusters (**Figure 2o**). AMPAR density sharply fell at longer distances from PSD-95 clusters as expected, consistent with the previous reports that surface AMPARs are highly enriched in synapses and are sparsely dispersed in the extra-synaptic sites ^1,5,11^. Therefore, the high-density regions, i.e., clusters of AMPARs identified earlier using DBSCAN, can be defined as synaptic clusters.

### 2.2. Differences in native surface AMPAR distribution in hippocampus vs cortex

One advantage of imaging native surface AMPARs in brain slices, as opposed to dissociated cultures, is the ability to study brain-region specific AMPAR distribution and its regulation. We therefore compared the hippocampus to the cortex of Thy1-YFP mice at 6 months of age (**Figure 3a-e**). By dSTORM, the relative density of total AMPARs was determined by the total AMPAR localization density per field of view (FOV) including all neurons regardless of the presence of YFP. It was 1.9x *larger* in the hippocampus than in the cortex (**Figure 3f** and **Table S5**). This is consistent with previous histoblot ^41^ and high-resolution mass spectrometry ^42^ studies. However, using super-resolution and cluster analysis, we can accurately quantify and compare synaptic clustering of AMPARs. Across 3 mice, we imaged at least 6 FOVs per mouse for each brain region and detected on average over 1500 AMPAR clusters per mouse for each brain region. We find that the fraction of AMPARs in synaptic clusters is *lower* in the hippocampus than the cortex by 2.9x (**Figure 3g**). Each datapoint on the plot represents the average measured value per mouse. We find a similar trend when limiting the analysis to YFP-labeled pyramidal neurons, where the fraction of AMPARs in synaptic clusters is 2.5x lower in the hippocampus than the cortex (**Figure 3h**).

**Figure 3.**
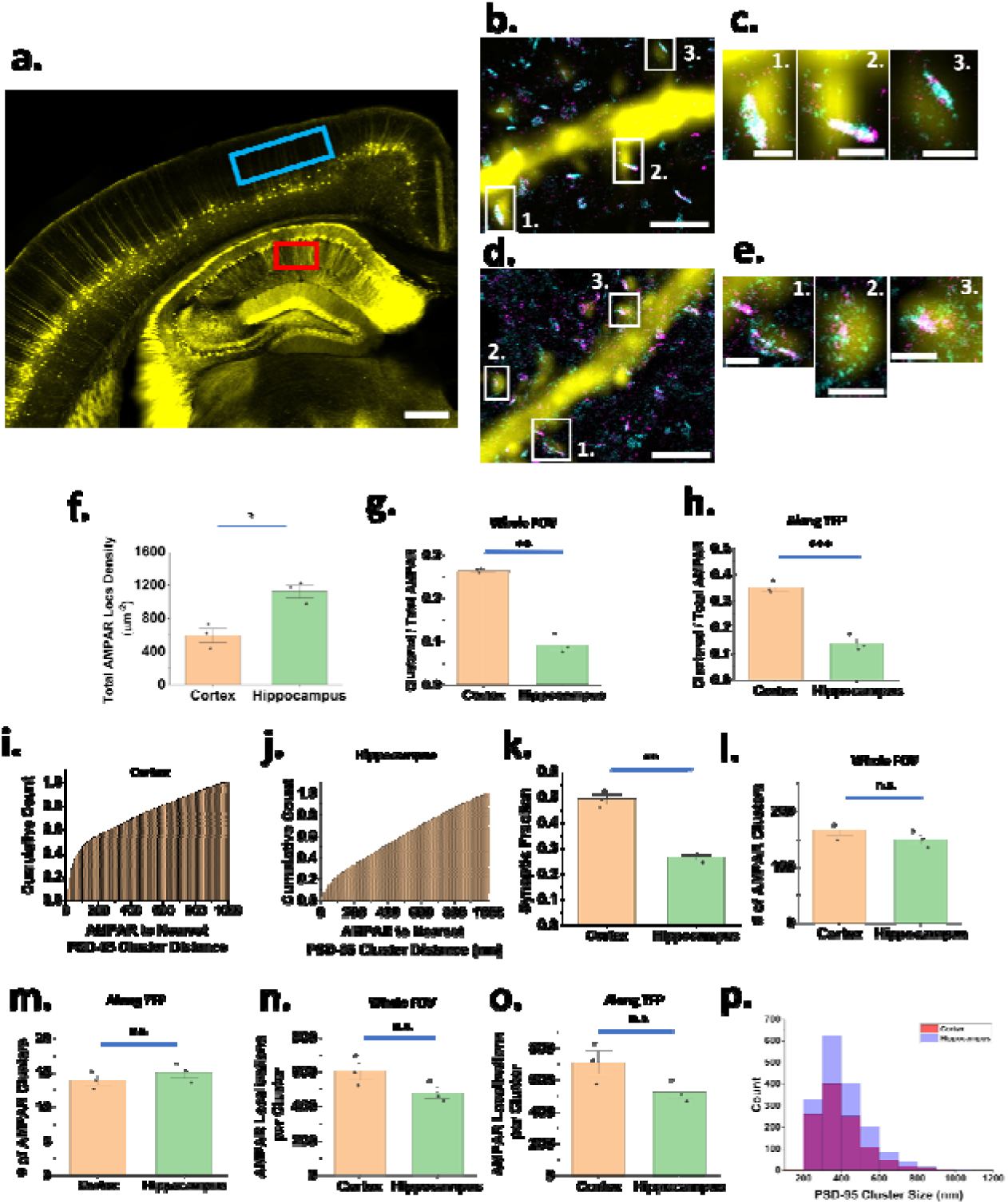
AMPAR distribution in hippocampus vs Cortex. **a,** Thy1-YFP mouse brain coronal section image using confocal microscopy. Super-resolution dSTORM imaging was performed in the regions highlighted for cortex (blue box) and hippocampus (red box) (Scale bar: 400 µm). **b,** Surface AMPAR distribution (cyan) along a YFP neuron (yellow) in the cortex of a 6-months old Thy1-YFP-H mouse, along with PSD-95 (magenta). (Scale bar: 2 µm) **c,** Zoomed in ROIs from b (Scale bar: 0.5 µm). **d,** Surface AMPAR distribution (cyan) along a YFP neuron (yellow) in the CA1 region of the hippocampus of a 6-months old Thy1-YFP-H mouse, along with PSD-95 (magenta). (Scale bar: 2 µm) **e,** Zoomed in ROIs from d (Scale bar: 0.5 µm). **f,** Total AMPAR localization density per imaging field of view (FoV) (p=0.042). **g,** Fraction of AMPARs in the whole FoV found in clusters (p=0.004). **h,** Fraction of AMPARs along a YFP neuron found in clusters (p=0.0004). **i,** Cumulative frequency distribution of the distance between each AMPAR localization and its nearest PSD-95 cluster in the cortex. **j,** Cumulative frequency distribution of the distance between each AMPAR localization and its nearest PSD-95 cluster in the hippocampus. **k,** Fraction of AMPARs lying within 140 nm of a PSD-95 cluster (p=0.003). **l,** Number of AMPAR clusters in the whole FoV (p=0.441). **m,** Number of AMPAR clusters on a YFP neuron (p=0.506). **n,** AMPAR localizations per cluster in the whole FoV (p=0.263). **o,** AMPAR localizations per cluster on a YFP neuron (p=0.226). p, Histograms of PSD-95 cluster size distribution for cortex (red) and hippocampus (blue). All error bars depict SE. For f-h and l-o: n=3 mice, at least 6 FoVs per mouse per brain region. Each point represents average value per mouse. For i,j,p: Averaged over 3 mice, at least 3 FoVs per mouse. For k: n=3 mice, at least 3 FoVs per mouse per brain region. Each point represents average value per mouse. ***:P<0.001, as determined using paired sample t-test; **: p<0.01, as determined using paired sample t-test; *: p<0.05, as determined using paired sample t-test; n.s.: p>0.05 as determined using paired sample t-test

We also tested whether CAM2 concentration played any role in the clustering differences observed. We increased the labeling of CAM2-Alexa 647 concentration from 2 µM to 5 µM in acute brain slices, and intracranially injected CAM2-Alexa 647 (2 µL, 50 µM) near the CA1 region of the hippocampus in mouse ^30^. In both cases, we observed the same lower AMPAR clustering in the hippocampus compared to the cortex, where it was 2.5x lower for 5 µM CAM2-AF647 and 4x lower for intracranially injected CAM2-AF647 (**Figure S6a-c**, **Table 6, 7**). Notably, the trend in AMPAR clustering was found to be reversed in primary (rat) neuronal cultures prepared from the hippocampus and the cortex (DIV 16) (**Figure S6d, Table S8**), where a 1.5x *increase* was observed in the hippocampal cultures compared to cortical cultures. Hence, the observed difference in AMPAR synaptic clustering between the hippocampus and the cortex in brain slices is likely independent of the concentration of AMPAR labeling and is different from the trend observed in dissociated primary neuronal culture.

Independent of AMPAR cluster analysis, the difference in synaptic distribution between the hippocampus and cortex is also suggested by measuring the AMPAR distance distribution around PSD-95 clusters along a YFP-positive neuron. There is a change in the slope of the cumulative distribution of AMPAR localizations at ∼100-140 nm in the cortex (**Figure 3i**, **Figure S7a**) and in the hippocampus (**Figure 3j**, **Figure S7b**). We therefore define this region around a PSD-95 cluster as the synaptic region. The fraction of AMPAR within this 140 nm synaptic region is 1.8x lower for hippocampus compared to the cortex (**Figure 3k**). We note that the exact definition of the synaptic region (for example, 100 nm vs 140 nm) doesn’t affect the observed trend (**Figure S7c, Table S9**). Combining the increased extrasynaptic fraction with increased number of total AMPAR in the hippocampus (**Figure 3f**), this means that the number of *extra-synaptic* AMPAR (= total number*extrasynaptic fraction) is higher in the hippocampus compared to the cortex.

We also measured the number of synaptic clusters over the FOV or along a YFP-positive neuron as well as the number of AMPAR within each cluster. We find no change between the hippocampus and the cortex for either of these measurements (**Figure 3l-o**). This suggests that the number and distribution of surface AMPAR within synaptic regions are the same for the two regions. It should be noted that clustered AMPARs are synaptic but synaptic does not mean clustered. Therefore, the additional AMPARs in the hippocampus are primarily located in the (sparse) extrasynaptic sites.We also compared the PSD-95 cluster size distribution and did not see a significant difference between cortex and hippocampus (**Figure 3p**).

Additionally in the hippocampus, we also tried to see if AMPAR clustering changes as a function of the distance from the soma layer. We did not see a significant difference up to 700 µm distance from the soma layer (**Figure S8**).

### 2.3. Changes in the Native surface AMPAR distribution at the onset of tauopathy

Another important advantage of imaging native surface AMPARs in brain slices is the ability to study their changes and therefore synaptic strength as a function of pathology in adult mice. Here, we focus on tauopathy, which is a leading cause of dementia-related illnesses such as Alzheimer’s disease.^43^ We use transgenic PS19 mice that model tauopathy by expressing human tau with P301S mutation associated with the frontotemporal dementia. This mutation induces accelerated pathology including tau hyperphosphorylation, formation of neurofibrillary tangles, synaptic deficits neurodegeneration, and cognitive decline. We crossbred Thy1-YFP-H mice with PS19 mice, such that the resulting PS19 mice (referred to as “PS19”) also expresses YFP in a subset of forebrain excitatory pyramidal neurons. At 6 months of age, there is no apparent loss of neurons in the double mutant PS19:Thy1-YFP-H mice compared to their Thy1-YFP-H only littermates (referred to as “WT”), but deficits in both long-term potentiation of excitatory synaptic strength and cognition have been reported via electrophysiology and behavioral studies. ^36,44^

To shed more light on the effect of tauopathy on synaptic strength, we observed how surface AMPAR organization changes in these mice. As seen in **Figure 4a** and **4b** showing WT and PS19 hippocampus, the difference is not obvious; nevertheless, subtle and important changes are present, which are revealed upon further quantitative analysis. The relative localization density of total AMPAR (as previously described for Figure 3f) is 1.6x reduced in the hippocampus of PS19 compared to the WT (**Figure 4c**, p = 0.023 and **Table S10**). This reduction in AMPAR expression have been observed previously in immunohistological studies (which included both surface as well as internal AMPAR)^36,41^ and immunoblots of membrane fractions (which included surface AMPAR).^41^ The relative localization density of total AMPAR in the cortex is also reduced by 1.7x, although the larger variance between mice makes this not statistically significant (Figure 4c, p=0.084).

**Figure 4.**
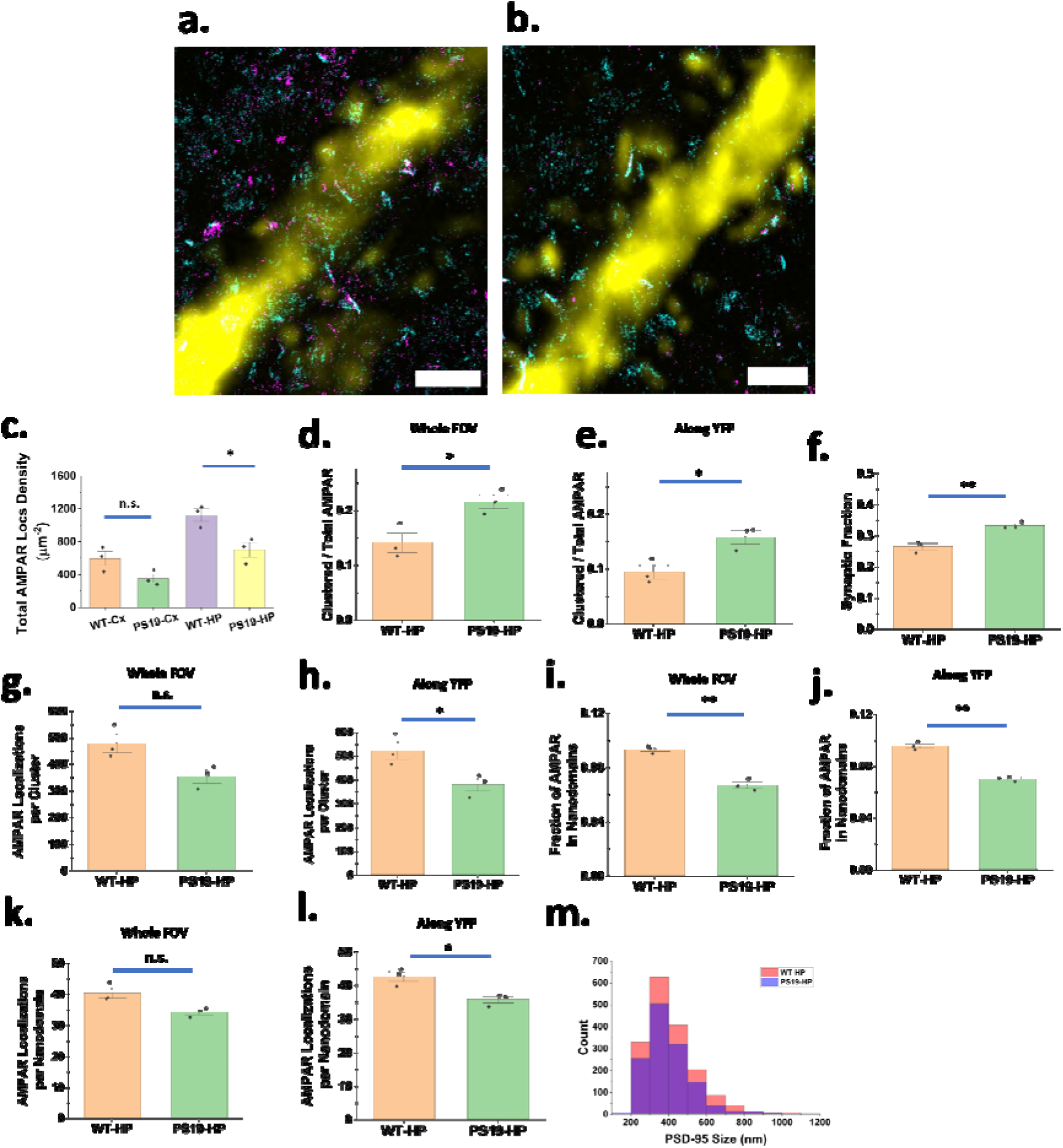
AMPAR organization in PS19 mice. Surface AMPAR distribution (cyan) along a YFP neuron (yellow), along with PSD-95 (magenta) in the hippocampus of a 6-months old **a,** WT and **b,** PS19 mouse (Scale bar: 2 µm). **c,** Total AMPAR localization density per imaging FoV (p=0.084, p=0.023). **d,** Fraction of AMPARs in the whole FoV found in clusters (p=0.024). **e,** Fraction of AMPARs on a YFP neuron found in clusters (p=0.035). **f,** Fraction of AMPARs lying within 140 nm of a PSD-95 cluster (p=0.009). **g,** AMPAR localizations per cluster over whole FOV (p=0.056). **h,** AMPAR localizations per cluster along a YFP neuron (p=0.048). **i,** Fraction of AMPARs in a cluster over the whole FoV found in nanodomains (p=0.001). **j,** Fraction of AMPARs in a cluster on a YFP neuron found in nanodomains (p=0.002). **k,** Number of AMPARs per nanodomain over the whole FoV (p=0.051). **l,** Number of AMPARs per nanodomain in a YFP neuron (p=0.027). m, Histograms of PSD-95 cluster size distribution for WT hippocampus (red) and PS19 hippocampus (blue). All error bars depict SE. For c-e and g-l: n=3 mice each for WT and PS19, at least 6 FoVs per mouse per brain region. Each point represents average value per mouse. For f, m: n=3 mice each for WT and PS19, at least 3 FoVs per mouse. Each point represents average value per mouse. **: p<0.01, as determined using two-sample t-test (Welch Correction); *p<0.05, as determined using two-sample t-test (Welch Correction); n.s.: p>0.05 as determined using two-sample t-test (Welch Correction).

Next, we wanted to observe possible changes in synaptic clustering of AMPARs. However, the decreased number of total AMPAR localizations in the PS19 hippocampus may indicate that the AMPAR clusters are simply undetectable because of their lower density with the DBSCAN parameters used. Indeed, upon visual inspection, we found relatively high-density regions not classified as AMPAR clusters with the DBSCAN parameters used for WT mice (Extended Data Figure 9). Therefore, we reduced our DBSCAN parameters by 1.6x for the PS19 hippocampus data based on the ratio of average AMPAR localizations per field of view detected in the hippocampus of WT and PS19 mice (for WT, the parameters are unchanged). We find that the fraction of AMPARs in synaptic clusters (either across the whole FOV or along a YFP neuron) is *higher* in the hippocampus of PS19 than in the hippocampus of WT (**Figure 4d**, p=0.02; **4e**, p=0.03 and **Table S11**).

To verify whether the fraction of synaptic AMPAR is indeed increasing in the PS19 hippocampus compared to WT, independent of DBSCAN parameter scaling, we compared the synaptic fraction of AMPARs (within 140 nm of PSD-95 cluster) along a YFP-positive neuron. We observed a significant increase in synaptic fraction (**Figure 4f**, p=0.008) consistent with the observation of higher fraction of clustered AMPARs in the PS19 hippocampus (Figure 4d, e). This suggests that tauopathy is associated with an increase in synaptic fraction and a decrease in extrasynaptic fraction of surface AMPARs in the hippocampus.

Given that the PS19 hippocampus has a lower total of number of AMPAR per field of view compared to the WT hippocampus (Figure 4c), the question is where is this decrease coming from? This decrease could be happening due to a decrease in the number of synaptic and/or extrasynaptic AMPAR. If both were decreasing by the same amount, the fractional change would be the same. However, since extrasynaptic AMPAR fraction is decreasing while the fraction of AMPARs in synaptic clusters is increasing (Figure 4d, e), we propose that the decrease in total number of AMPAR in the PS19 hippocampus is primarily due to a loss in extrasynaptic AMPAR. This agrees with previously reported decrease in extrasynaptic AMPARs in the PS19 hippocampus observed at 10 months of age using SDS-digested Freeze Fractured Replica Labeling (SDS-FRL).^41^

As in Figure 3l and 3m, we used cluster analysis to compare the number of AMPAR localizations per synaptic cluster for the hippocampi in WT and PS19 mice. There is a statistically significant difference for clusters in the YFP-labeled neurons (**Figure 4h**, p=0.048) but not for all the clusters in the FOV (**Figure 4g**, p=0.055). This suggests that the number of AMPARs per synaptic cluster on YFP-positive neurons is likely to be decreasing by a factor of 1.4 in the PS19 hippocampus.

To probe the AMPAR organization within a synaptic cluster, we measured the fraction of AMPARs found in nanodomains over the whole FOV and along YFP neurons. We find that the fraction is lower in the PS19 hippocampus compared to the WT hippocampus (**Figure 4i**, p=0.001; 4j, p=0.002). In addition, the number of AMPAR localization per nanodomain is significantly lower along a YFP neuron (**Figure 4l**, p=0.027) but not in the whole FoV (**Figure 4k**, p=0.051). These results together suggest that the organization of AMPARs in synaptic nanodomains in the PS19 hippocampus is likely disrupted: that is, AMPARs are not as well-organized in synaptic nanodomains in PS19 hippocampus, as compared to WT hippocampus.

The PSD-95 size distribution did not show a significant difference between PS19 hippocampus and WT hippocampus (**Figure 4m**). Additionally, we did not observe any significant changes to the AMPAR distribution in the PS-19 cortex compared to the WT-cortex (**Figure S10**).

## 2. Discussion

A substantial body of evidence is present in existing literature that associates the diffusion and nanoscale organization of AMPAR in the post-synaptic membrane of neurons as key indicators of synaptic strength. Changes in their organization and other biophysical properties form the underlying mechanism behind various forms of synaptic plasticity. Despite this significance, super-resolution imaging of native, surface AMPARs directly in adult brain tissue, particularly to examine differences across brain regions or disease states, has remained limited. To address this gap, we used the CAM2 probe to label native surface AMPARs in 6-month-old Thy1-YFP-H mouse brain tissue (Figure 1a). Combining dSTORM imaging with refractive-index matching, adaptive optics, multi-step drift correction, and INSPR enabled visualization of CAM2-labeled AMPARs alongside PSD proteins such as PSD-95 with sub-10-nm lateral and sub-30-nm axial localization precision (Figure 1b, 2a-j, Figure S2).

To test our methodology, we measured certain properties that have been previously reported in the literature such as the distance of PSD-95 and Homer-1 clusters from the surface AMPAR, correlation of PSD-95 and AMPAR clusters that form part of a trans-synaptic nanocolumn as well as the size of AMPAR nanodomains (Figure 2k-o). We found excellent agreement. Additionally, we compared PSD-95 labeling with and without CAM2-labeling step and in the presence of a conventional anti-GluA2/3 antibody to ensure that PSD-95 labeling is not affected by CAM2-labeling (Figure S4, S5).

In order to examine the expression of surface AMPAR as a function of brain region, we compared the CA1 region of the hippocampus with the motor and somatosensory cortical regions that lie directly above the hippocampus in coronal sections of 6-month-old Thy1-YFP-H mice brains (Figure 3a-e). While it is known that the amount and subunit composition of total AMPARs are different between these two regions^42^, a quantification of their differences in nanoscale synaptic distribution is lacking. Despite the proximity of these two regions, we found a significant difference in the synaptic distribution of surface AMPARs. Specifically, we found over two-fold decrease in the proportion of AMPARs organized in synaptic clusters in the CA1 region of the hippocampus compared to the motor and somatosensory cortex (Figure 3g). This trend was robust across labeling conditions, including analysis restricted to YFP-positive neurons (Figure 3h), increased CAM2 concentration in acute slices, and intracranial CAM2 delivery in vivo (Figure S6). Moreover, neither the number of AMPAR clusters per field of view nor the number of receptors per cluster differed significantly between regions (Figure 3l-o).

Independent of AMPAR clustering analysis, we quantified the synaptic and extrasynaptic organization of AMPAR by measuring the distance of AMPAR localizations around a PSD-95 cluster (Figure 3 i, j). This revealed a decrease in the synaptic fraction and thus, a corresponding increase in the extra-synaptic fraction in the CA1 region of the hippocampus compared to the motor and somatosensory cortex (Figure 3k). We recognize the potential bias towards stronger PSD-95 clusters in our synaptic fraction measurement results, since we found only a third of the YFP spines contained the PSD-95 clusters as defined by our conservative clustering parameters (Methods). For the AMPAR synaptic fraction measurements, we identified the PSD-95 cluster first and then identified all AMPAR localizations within 1 micron distance. However, given that the same methodology was used for synaptic fraction measurements in both hippocampus and cortex, the observed differences still remain valid. Taken together with the overall greater abundance of surface AMPAR observed in the hippocampus (Figure 3f), these results imply that the hippocampus contains a substantially larger extrasynaptic AMPAR pool than the cortex.

Extrasynaptic AMPARs are thought to represent a highly mobile surface reserve pool that can be rapidly recruited into synapses during the early expression of synaptic potentiation through activity-dependent diffusion trapping and synaptic stabilization mechanisms.^45,46^ Importantly, synaptic plasticity, including LTP, is robust in both hippocampus and the cortex. However, hippocampal and cortical networks are proposed to play complementary roles in memory processing across timescales, with hippocampal circuits supporting rapid encoding and associative plasticity, while cortical regions contribute to the longer-term consolidation, stabilization, and organization of learned representations^34,35^. In this context, the increased extrasynaptic AMPAR fraction observed in the hippocampus may reflect a larger pool of receptors available for rapid synaptic recruitment during activity-dependent plasticity, potentially aligning with the specialized role of hippocampal circuits in memory formation.

We also examined changes in the nanoscale organization of surface AMPARs in a pathological mouse model. Specifically, we observed changes in the hippocampus of a PS19 mouse model of tauopathy at 6 months of age. This time-point was chosen as it precedes visible neurodegeneration or brain atrophy but coincides with the onset of synaptic and learning deficits.^36^ While previous studies have reported a disruption of AMPAR trafficking pathways or a gross-level loss of synaptic and extra-synaptic AMPARs in Alzheimer’s disease mouse models that display tau burden and amyloid beta accumulation,^41,47^ we observed a more complex change in its distribution (Figure 4). We found that while the total number of AMPARs (comprised of synaptic and extra-synaptic AMPARs) decreased in the tauopathy model (Figure 4c), *the fraction* of AMPARs located in synapses is in fact higher (Figure 4d-f). Consequently, the decrease in total number of AMPARs primarily comes from the extra-synaptic (as opposed to synaptic) population. Given the role of extrasynaptic AMPARs in receptor exchange and plasticity, this redistribution is likely to constrain synaptic adaptability. Within synapses, we also detect a decrease in the proportion of AMPARs organized in synaptic nanodomains (Figure 4i,j) along with a modest decrease in the number of AMPARs per nanodomain as well as per synaptic cluster (Figure 4k,l). Given that the disruption in the nanoscale organization of synaptic AMPARs has been shown to affect the synaptic transmission,^10,48,49^ this points to a further synaptic weakening and deficient neurotransmission in the PS19 hippocampus.

Direct quantal recordings in PS19 hippocampus remain relatively limited; however, available whole-cell studies demonstrate that mEPSC amplitude is altered in CA1 pyramidal neurons, indicating quantal-level synaptic dysfunction beyond circuit-level field measures.^50^ Importantly, hippocampal plasticity phenotypes in PS19 mice are stage- and protocol-dependent: Bold et al.^50^ report reduced basal synaptic strength together with aberrantly enhanced theta-burst–induced LTP at 16–18 weeks of age, which they attribute to early network disinhibition, whereas Yoshiyama et al.^36^ report impaired LTP at later stages at ∼6 months of age associated with more advanced synaptic degeneration. Thus, tauopathy produces dysregulated synaptic function rather than a uniform change in plasticity magnitude.

Taken together, our nanoscale measurements identify a mechanistically meaningful subsynaptic substrate for synaptic dysfunction in tauopathy. By resolving both synaptic nanodomain disruption, which may directly affect quantal efficacy, and extrasynaptic AMPAR depletion, which constrains receptor exchange during synaptic plasticity, our data provide a structural framework that helps explain weakened basal transmission and dysregulated plasticity. The AMPAR nanodomain disruption can be an early indicator of the eventual loss of spines and neurons in PS19 mice at a later age. ^36,51,52^ Our results also help provide a possible molecular-level explanation for the learning deficit in tauopathy models.^53^

As these nanoscale changes in tauopathy are not readily accessible using conventional lower-resolution imaging approaches, our method has the potential to reveal previously unresolved alterations in AMPAR organization across other disease models associated with cognitive deficits. Future improvements in dSTORM imaging throughput, together with learning-based analysis approaches, will further enable larger-scale extraction of synaptic and nanoscale organizational features from thick tissue samples. Estimating the absolute protein numbers from the single-molecule blinking data would also be beneficial in determining the absolute AMPAR loss during tauopathy. To avoid large uncertainties in such an estimation will require using an appropriate photophysical model and a calibration/training dataset that is carefully matched with the experimental conditions in terms of local environment, imaging buffer and labeling density^54,55^.

Several important directions remain for future work. First, multiplexed imaging of surface AMPARs together with pathological tau would help determine whether local tau burden is spatially associated with synapse-specific AMPAR nanoscale disorganization. Second, subunit-specific AMPAR imaging could clarify whether disease-associated changes involve the total surface AMPAR population or preferentially affect receptors with specific GluA-subunit compositions. Another important avenue for future work would be to identify the exact mechanistic explanations of the observed AMPAR dysregulation due to tauopathy. Although we did not observe a significant reduction in PSD-95 synaptic enrichment, preserved PSD-95 labeling does not exclude more specific tau-dependent changes in postsynaptic molecular coupling.^56^ It would also be interesting to study the effect on the synaptic enrichment of other scaffolding proteins such as SAP97^57^ and TARPs, in particular Stargazin^58,59^, that can affect AMPAR organization and trafficking. Tau–PACSIN1 interactions have also been implicated in the regulation of AMPAR trafficking and extrasynaptic AMPAR function.^60^ Identifying the exact mechanistic explanations would also help develop a therapeutic intervention that can restore the observed AMPAR dysregulation and potentially mitigate the synaptic dysfunction observed in tauopathies.

## 3. Conclusion

In this work, we observed the nanoscale organization of native, surface AMPARs for the first time in adult mouse brain slices using a combination of CAM2 labeling and dSTORM imaging strategies. We were able to observe the synaptic clustering of AMPARs colocalized with the PSD proteins as well as synaptic nanodomains.

We analyzed AMPAR distributions as a function of the brain region, comparing the hippocampus and the neighboring motor and somatosensory cortex. For WT mice, we found a smaller fraction of AMPARs in synaptic clusters in the hippocampus compared to the cortex, despite having a higher AMPAR density in the hippocampus. We furthermore showed that the additional AMPARs found in the hippocampus were primarily located in extra-synaptic sites.

The differences in AMPAR distribution might be related to different functions of the two brain regions in learning and memory.

In PS19 mice, we found a loss of balance in the proportion of AMPARs located inside versus outside synapses. We further found a reduction of AMPARs per synaptic cluster, combined with the disruption of their organization in nanodomain in PS19 mice. In the future, our method can also be applied to investigate nanoscale alterations in the distribution of surface AMPARs in mouse models of other pathological diseases or learning paradigms.

## 4. Experimental Section/Methods

### 4.1 Animals

All animals were housed, euthanized and dissected in accordance with the guidelines from the Institutional Animal Care and Use Committee at the University of Illinois at Urbana-Champaign (IACUC protocol #24039). Thy1-YFP-H mice were purchased from Jackson Labs (strain #003782). PS19 mice were also purchased from Jackson Labs (strain #008169). The mouse lines were maintained in-house by breeding the respective hemizygote transgenic mice with their noncarrier counterparts. The crossbreed Thy1-YFP x PS19 mouse line was created by breeding hemizygote male mice of one strain with hemizygote female mice of the other.

Genotyping was performed by Polymerase Chain Reaction from tail biopsies. The protocol and primers were used as per Jackson Labs instructions. Thy1-YFP-H mice used in this study were littermate controls of PS19:Thy1-YFP-H mice. For each genotype, three 6-month-old mice were used, with each group containing two males and one female.

Primary hippocampal and cortical cultures were prepared from E18 rat embryos obtained from timed-pregnant rats (Charles River).

### 4.2 Acute Slice Preparation and CAM2 staining

Mice were sedated using isoflurane and decapitated. Brains were then extracted within 5 min of decapitation and mounted onto a Leica vibratome. During sectioning, brains were kept in high osmolarity, ice cold cutting solution (2.5 mM KCl, 2.5 mM MgCl_2_.6H_2_O, 0.5 mM CaCl_2_.2H_2_O, 1mM NaH_2_PO_4_.H_2_O, 220 mM sucrose, 25 mM NaHCO_3_ and 20 mM glucose (dextrose) in ddH_2_O) with continuous bubbling of 95% O_2_/5%CO_2_ gas mixture. 200 µm acute slices of the brain were made using the vibratome and incubated in artificial cerebrospinal fluid (ACSF) (126 mM NaCl, 3 mM KCl, 2 mM MgSO_4_.7H_2_O, 2 mM CaCl_2_.H_2_O, 1mM NaH_2_PO_4_.H_2_O, 25 mM NaHCO_3_ and 14 mM glucose (dextrose) in ddH_2_O) for 30 min at 37°C under continuous bubbling of 95% O_2_/5%CO_2_ gas mixture to stabilize the brain slices. The slices were then incubated with 2 µM CAM2-Alexa Fluor 647 in ACSF for 2 hours at room temperature under continuous bubbling of 95% O_2_/5%CO_2_ gas mixture (or 5 µM for experiment with increased CAM2-Alexa Fluor 647 concentration). After washing 3x with ACSF, the slices were fixed with 4% PFA in PBS at 4°C for 24 hrs. After fixation, the slices were washed 3x with PBS and stored in PBS for cryosectioning and further labeling.

### 4.3 Stereotaxic surgery for intracranial injection of CAM2-AF647

Three-month-old Thy1-YFP mouse was anaesthetized with a mixture of ketamine (100 mg/kg) and xylazine (10 mg/kg). When deeply anaesthetized, the mouse was placed on a heating pad to maintain body temperature during surgery. Maintenance anesthesia was given at 1% isoflurane with O2. Complete anesthesia was confirmed by tail pinch, and the head was fixed in a stereotaxic apparatus. Using a Hamilton syringe, the hippocampus (A/P, −2.5 mm from bregma; L, −2.0 mm; D/V, −1.8 mm) received a left hemisphere stereotaxic injection of 2.0 μl brain extract, at a speed of 1.0 μl/min, for CAM2-AF647 (50 uM). Following injection, the needle was kept in place for an additional 1 min before gentle withdrawal to minimize reflux^61^. For the same mouse, the right cerebral cortex/white matter (bregma: AP −2.50 mm, ML +2.00 mm; dural surface: DV −0.80 mm)^62^ received stereotaxic injection as well of of 2.0 μl brain extract, at a speed of 1.0 μl/min, for CAM2-AF647 (50 uM).

The surgery was completed by suturing the scalp and allowing the mouse to recover with antibiotics (Neosporin) and analgesic treatment (Lidocaine and Prilocaine Cream). Mice were monitored until recovery from an aesthesia and checked following surgery. All animal experiments described here were approved by the animal care and use committee of University of Illinois at Urbana-Champaign.

### 4.4 Antibody-dye conjugation

For goat anti-rabbit secondary antibody conjugation with CF568 dye, Antibody (in PBS) is mixed in equal parts with NaHCO3 buffer (fresh, 0.1 M, pH 8.3). Dye : antibody are mixed in 15 : 1 ratio. The dye+antibody mixture is kept on a shaker at room temperature for 2 hrs (wrapped in aluminum foil to prevent photobleaching). Zeba spin desalting column (7K MWCO) is used to separate out free dye. For buffer exchange, Amicon Ultra-0.5 centrifugal filter (50 kD) is used to resuspend the antibodies back in PBS. (14000xg, 10 min, repeated three times).

### 4.5 Nanobody-dye conjugation

FluoTag®-X2 anti-PSD95 nanobody (#N3702-250ug, NanoTag Biotechnologies) was first conjugated to CF®568 maleimide (#92024, Biotium), according to the manufacturer’s protocol. The maleimide-conjugated nanobody went through Zeba desalting column (#89882, Thermo Scientific) to get rid of the excess maleimide dye, and was subsequently conjugated to CF®568 succinimidyl ester (#92131, Biotium) via NHS reaction to increase the number of dyes per nanobody. The final product was purified using Zeba desalting column (#89882, Thermo Scientific) first, and then with Amicon Ultra Centrifugal Filter (#UFC5005, Millipore Sigma).

### 4.6 Immunohistochemistry

CAM2 labeled 200 µm acute brain slices were embedded in optimal cutting temperature (OCT) (#12351753, Fisher Scientific) in cryosectioning molds (#27183, Ted Pella). At this point, slices were either immediately cryo-sectioned or stored at −80°C for long-term storage. Gelatin-subbed coverslips were prepared for collecting cryosections. Coverslips (#64-0715, Warner Instruments) were first sonicated with 1M KOH for 15 min, followed by sonication for 15 min with 95% ethanol. Coverslips were then dipped in warm (∼50°C), filtered 0.5% gelatin and 0.05% chromium potassium sulphate solution in ddH_2_O for 1 min and were dried in an oven at 50°C overnight. 30 µm cryosections of the OCT embedded 200 µm acute slices were made using a Leica cryostat (CM3050 S) and collected on the gelatin-subbed coverslips.

Sections at either edge of the 200 µm slice were discarded. The 30 µm sections were then incubated in the blocking buffer comprised of 0.2% Triton-X 100 (#648466, Millipore Sigma) and 5% Normal Goat Serum (NGS) (#005-000-121, Jackson Immunoresearch) in PBS for 12 hrs at room temperature on a shaker. The solution was then replaced with either 33 nM unconjugated anti-Homer primary antibody (#16003, Synaptic Systems) or 33 nM unconjugated anti-GluA2/3 primary antibody (AB1506, Sigma-Aldrich) or 33 nM anti-PSD95 nanobody conjugated to CF568 dye in blocking buffer for 24 hrs at room temperature (RT) on a shaker. The slices were then washed 3x with PBS, with 20 min incubation at RT on a shaker between each wash. After the anti-Homer primary antibody staining, slices were incubated with 33 nM goat anti-rabbit secondary antibody (#ab7085, Abcam) conjugated to CF568 in blocking buffer for 24 hrs at RT on a shaker. After the anti-GluA2/3 primary antibody staining, slices were incubated with 33 nM goat anti-rabbit secondary nanobody pre-conjugated to Alexa Fluor 647 (#N2402-AF647-S, NanoTag) in blocking buffer for 24 hrs at RT on a shaker. The slices were then washed 3x with PBS, with 20 min incubation at RT on a shaker between each wash. Afterwards, the slices were postfixed with 4% PFA in PBS for 2 hrs at RT on a shaker. PFA was then washed out three times with PBS.

### 4.7 Primary neuron culture and CAM2 staining

Timed pregnant rats were euthanized in a CO_2_ chamber and embryos were extracted from them after decapitation. Hippocampuses (or cortices) were dissected out from the brains of these embryos after removing the meninge. Hippocampal (or cortical) neurons were dissociated and cultured as described previously.^11^ At 13 days in vitro (DIV), cells were transfected with 1 µg GFP plasmid (pEGFP-C1, Clontech, Genbank Accession #: U55763, https://www.addgene.org/vector-database/2487/) using 4 µg Lipofectamine 2000 (Invitrogen, Cat. #11668019). After 48 hours, coverslips with transfected primary hippocampal (or cortical) neurons were then incubated in 1 mL culture media with 1µM CAM2-Alexa 647 for 4 hrs at 37°C. After this, coverslips were washed thrice with 1 mL culture media and fixed with a solution containing 4% PFA and 4% sucrose in PBS for 20 min at room temperature (RT). PFA was washed out with 1 mL PBS thrice. Neurons were then incubated in the blocking buffer comprised of 0.2% Triton-X 100 (#648466, Millipore Sigma) and 5% Normal Goat Serum (NGS) (#005-000-121, Jackson Immunoresearch) in PBS for 20 min at room temperature on a shaker. Neurons were then incubated with 12 nM anti-PSD95 nanobody conjugated to CF568 dye in blocking buffer for 1 hr at RT on a shaker. The nanobody was then washed out three times with PBS. Afterwards, the neurons were postfixed with 4% PFA in PBS for 15 min at RT on a shaker. PFA was then washed out three times with PBS.

### 4.8 Airyscan Confocal Imaging

For Airyscan confocal imaging, Zeiss LSM900 microscope was used with either 63x or 10x microscope objectives. Zen Blue (Zeiss) software was used for image acquisition.

### 4.9 Microscope Setup for STORM Imaging

The custom microscope setup was built around an Olympus IX-71 microscope body. Four lasers were used to illuminate the sample at different wavelengths: 639 nm (Coherent Genesis MX639-1000 STM), 561 nm (Coherent Genesis MX561-500 STM), 488 nm (Coherent Saphire), 405 nm (Coherent OBIS). All excitation beams were magnified by 20-30x and only the central portion was collected using an aperture to get a uniform illumination across the field of view. A Nikon 100x NA1.35 silicone oil immersion objective was used (CFI SR HP Plan Apochromat Lambda S 100X Sil) for illuminating the sample and collecting the fluorescence emitted. A quad-band dichroic mirror (ZT 405-488-561-640RPC, Chroma) was used to separate excitation from emission. A 700nm/75nm bandpass emission filter (ET700/75m, Chroma) was used for Alexa fluor 647. A 600nm/75nm bandpass emission filter (Chroma) was used for CF568. A 525nm/50nm bandpass emission filter (Chroma) was used for Alexa fluor 488 and YFP. The images are collected using an EMCCD camera (ixon^+^ DU897, Andor). A deformable mirror (Multi-3.5-DM, Boston Micromachines) is placed at the pupil plane in the emission path to correct aberrations and introduce astigmatism for 3D dSTORM. This deformable mirror (DM) has 140 actuators that shape the reflective membrane surface depending on the voltage applied to them. The DM needs to be aligned carefully such that it sits precisely at the pupil plane with near normal incidence angle and the size of the pupil is only slightly less than the DM aperture. If the pupil is larger, DM will be overfilled, resulting in signal clipping. If the pupil is too small, DM will be underfilled resulting in underutilization of the DM actuators and poorer mirror mode generation. The size of the pupil at the back aperture of the microscope objective is given by 2*NA*f, where NA is the numerical aperture and f is the focal length of the objective. For our objective, the pupil size of 5.4 mm was diminished down to 3.75 mm using an array of lenses so that it is close to the 4.4 mm DM aperture. The angle of incidence at the DM was ∼19 degrees. The final magnification of the image at the EMCCD was 162x, resulting in 99 nm effective pixel size. An 850 nm LED was used to illuminate the sample from the top and the transmission images were collected using a separate CCD camera (DMK 23U74, The Imaging Source). The sample was placed on an xyz piezo stage (Mad City Labs) which was placed on a mechanical stage (ASI). The entire microscope setup was controlled using a custom LabVIEW code. A custom MATLAB code was used to generate the theoretical mirror modes for the DM,^63,64^ and the actuator voltage values for each mode were fed to the DM through the control LabVIEW code. Instrument-induced static aberrations in the system were corrected by cycling through the first 30 mirror modes and 11 amplitudes for each mode for a 100 nm sized fluorescent bead sample (T7279, Thermo Fisher) and finding a linear combination of each mode and its corresponding amplitude that maximizes the fluorescence intensity of the bead point spread function (PSF).

### 4.10 dSTORM Imaging

For dSTORM imaging, coverslips were mounted onto a holder (A7816, Thermo Fisher) along with the imaging buffer containing 10% Oxyfluor (OF-0005, Oxyrase Inc.) and 10 mM sodium DL-lactate (71720, Sigma-Aldrich) in PBS as the oxygen scavenging system. The imaging buffer also contained 100 mM Cysteamine/Mercaptoethylamine (M9768-5G, Sigma-Aldrich) as a reducing agent and 35% 2-2-thiodiethanol in PBS for refractive index matching. A plastic lid was placed on the holder with mounted coverslips and the imaging buffer and the chamber was sealed using black aluminum foil tape (T205-1.0, Thorlabs). Before starting dSTORM imaging, first a ±1 µm z-stack with 100 nm step size above and below the plane of focus at the region of interest was collected in the IR (850 nm) transmission imaging CCD camera. This served as the reference stack for real-time drift correction. Then, a z-stack of YFP dendrites was collected using 488 nm laser excitation to compare colocalization with the rest of the channels later. Alexa fluor 647 signal was then bleached at 3-4 kW/cm^2^ 639 nm laser power density until single molecule blinking could be observed. During the bleaching step, the excitation beam angle was increased almost to epi mode to bleach most of the out-of-focus fluorescence background. The beam angle was then lowered such that it was just above the critical angle. Weak astigmatism was introduced in the system using the deformable mirror. 50,000 frames of dSTORM images at 50 ms EMCCD exposure time and 100 EM gain were collected for each field of view (FOV). Before imaging CF568, YFP was completely photobleached using 488 nm laser to avoid any YFP leakage into the CF568 channel. CF568 dSTORM imaging was performed using 3-4 kW/cm^2^ 561 nm laser power density in the same way as Alexa fluor 647, except 75,000 total frames were collected for CF568. Additionally for CF568 imaging, 405 nm laser was illuminated for 50 ms after every four CF568 acquisition frames. 405 nm laser intensity was gradually increased through the course of imaging from 10-200 W/cm^2^ to ensure a constant CF568 blinking rate.

### 4.11 Chromatic Aberration Correction

Despite the use of advanced microscope objectives and achromatic doublet lenses in the microscope setup, a small shift due to chromatic aberration can persist. When measuring nanoscale distances, it becomes important to accurately correct for the chromatic aberration shift between different colors. For this purpose, we used a silver-coated coverslip with 100 nm sized holes spaced at 1.5 µm distance away from each other. This nanohole patterned coverslip was used to map and register different channels to calculate the chromatic aberration shifts using a custom MATLAB code.^11^ Additionally, it is possible that the chromatic aberration is depth dependent.^65^ To test and measure this, tetraspeck beads (T7279, Thermo Fisher) with 100 nm diameter that fluoresce in 4 different channels (blue, green, orange, dark red) were embedded in 2% agarose gel and the chromatic aberration shift between green, orange, and dark red channels, corresponding to Thy1-YFP, CF568 and CAM2-Alexa Fluor 647 signals respectively, was measured as a function of depth (fig. S2g). While the difference in lateral chromatic shift was not large up to 30 µm imaging depth, axial chromatic shift was observed. This shift was corrected in the Thy1-YFP z-stacks and CF568 localization data with reference to the Alexa Fluor 647 channel.

### 4.12 Automated Dendritic Spine Detection and Segmentation

Conventional spine detection and modeling software such as Imaris and Neurolucida did not work well for the diffraction limited YFP images collected in the brain slices using the widefield microscope due to a poorer signal to noise ratio compared to dissociated neurons or confocal imaging even after rolling ball background subtraction (radius = 10 pixels) in FIJI ^66^. Hence, a recently developed machine learning based algorithm called deepd3 was used for automatically detecting dendrites and spines.^67^ Spines and dendrites were detected in the background subtracted YFP z-stack images using deepd3 and the predictions were segmented using a recently developed segmentation algorithm Startdist to separate overlapping spines in 3D. ^68^ Stardist was originally developed for segmenting fluorescence images of cell nuclei, however we find that it works well for spines also. An additional filtering criterion was used to keep only those spines that are in focus within ± 500 nm of the focal plane when comparing colocalization with SMLM data as that is roughly the z-range for the astigmatism used. Detailed method is described separately.^69^

### 4.13 dSTORM Image Analysis

dSTORM images were analyzed using INSPR. Parameters used in the software for pupil estimation were as follows: Background subtraction (median filter) = ON, box size = 32, distance threshold = 26, initial threshold = 10, segmentation threshold = 15. Z-range = −1 to 1 µm with 0.1 µm increment, Z shift mode = OFF, Wyant Zernike order = 64, Bin_lowBound = 50, Iteration = 8, Min similarity = 0.6, OTFscale = 2. Localizations were reconstructed using ThunderSTORM plugin in FIJI.^70^ ThunderSTORM was also used for conventional single-molecule localization using elliptical Gaussian PSF when comparing INSPR analysis performance in Extended Data Fig. 2c-f. Since the real-time drift correction employed in the microscope could not correct for lateral (xy) drift below 1 pixel accuracy, an additional drift correction step using 2D cross-correlation was performed in ThunderSTORM plugin for correcting lateral drift with sub-pixel accuracy. Drift for CF568 data was further corrected with reference to the first frame of Alexa Fluor 647 data to avoid any drift between the two channels. Chromatic aberration was also corrected for the CF568 data with reference to Alexa Fluor 647 channel using nanohole registration as described earlier. The chromatic aberration shift in diffraction limited YFP images was corrected with reference to Alexa Fluor 647 channel using cross-correlation between the reference nanohole images for the two channels. Additional depth dependent axial chromatic shift (as described earlier) was applied to CF568 and YFP images with respect to Alexa Fluor 647 channel.

Custom-written Python codes were used to analyze the single-molecule localization data obtained from INSPR analysis. DBSCAN algorithm ^37^ plus a minimum requirement on localizations in clusters were used to detect clusters of AMPARs (min points: 120, distance: 120 nm, min requirement: 160) and Homer1/PSD-95 (min points: 90 for Homer1 / 30 for PSD-95, distance: 120 nm, min requirement: 120) in 3D. The DBSCAN parameters were reduced by 1.6x for AMPAR cluster detection in PS19 hippocampus data based on the ratio of average AMPAR localizations per field of view detected in the hippocampus of WT and PS19 mice. To account for difference in AMPAR density in each cluster, we determine the density threshold for nanodomains based on the density of the cluster. For this purpose, the number of other localizations within 30 nm of each localization in a cluster is determined. Then the mean and standard deviation of the number of localizations for the cluster is calculated. DBSCAN was used to identify nanodomains inside the cluster (min points: mean+1.5 standard deviation, distance: 30 nm, min requirement: 20). Concave hulls of clusters and nanodomains are built using alphashape (inverse of alpha: 500 nm for clusters, 250 nm for nanodomains).

Size of each nanodomains is calculated as twice the average distance between each localization to the center. For YFP specific analysis, clusters of AMPAR-CAM2-Alexa 647 and Homer1/PSD-95-CF568 are assigned to their nearest segmented dendrite and spines if they fall within a minimum distance threshold (50 nm to dendrite and 100 nm to spines). A new distance threshold is set for each spine as the maximum distance to clusters assigned to it. Non-cluster localizations within the distance thresholds are then assigned to the spines and dendrite.

For two-color synaptic fraction analysis, AMPAR localizations associated with the YFP neuron are assigned to their nearest PSD-95 clusters of the neuron, and their distances to the cluster are calculated to obtain the synaptic distance distribution of AMPAR. We optimized the clustering parameters to define PSD-95 localization clusters such that any low-density background in the field of view is not being counted as clusters and ensure the accuracy of AMPAR distance measurements from the well-defined PSD boundary. This has limitations in that some low density, sparser PSD-95 clusters in the spines would get excluded, especially when looking at single-molecule localizations as opposed to diffraction limited images. However, having false PSD-95 cluster detections would be more detrimental to the accuracy of the AMPAR distance measurements from PSD boundary, and hence we decided to err on the side of caution.

For the distance of AMPAR to PSD-95/Homer1, a minimum centroid to centroid distance of 100 nm first finds pairs of AMPAR and PSD-95/Homer clusters. The distance between the two clusters is then calculated along their third principal component analysis (PCA) component directions (direction they orient toward each other). The 3D paired cross-correlation function (PCF) between two clusters is calculated using the formula below.^13^

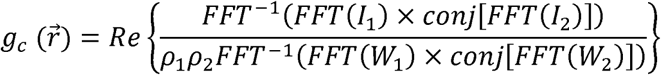

I_1_ and I_2_ are matrices of the AMPAR and PSD-95 localizations inside the cluster pairs (with minimum size requirement of 150 for each) in 3D with 20 (for these, need to check what works eventually) nm voxel size. W_1_ and W_2_ are shape functions of the clusters, which is 1 inside their concave hulls (inverse of alpha: 200 nm) and 0 outside, and ρ_1_ and ρ_2_ are the cluster densities. For simulated PCF data, uniformly distributed localizations are generated inside the concave hulls of the actual clusters with the same densities.

## Acknowledgements

We thank Fernando Rigal (UIUC) for his help in confocal and dSTORM imaging, respectively. We also thank Gregory Tracy (UIUC) and Ki H. Lim (UIUC) in Dr. Hee Jung Chung’s lab for their help in maintaining transgenic mice lines.

## Data Availability Statement

All data needed to evaluate the conclusions in the paper are present in the paper and/or the Supplementary Information. The raw dSTORM images acquired for this study run into several terabytes and hence it is not practical to upload onto a public repository. However, INSPR-analyzed files for every dSTORM experiment performed in this study, containing single-molecule localization information, are deposited at Dryad: https://doi.org/10.5061/dryad.xpnvx0kqc.

The raw .tiff files for all the dSTORM reconstructed images displayed in the paper as well as all the image analysis codes are also available there. All data used in this paper will be made freely available to those who request and provide a mechanism for feasible data transfers (such as physical hard disk drives or cloud storage).

