## Supplementary Tables and Figures for "Synaptic Organization of Surface AMPARs Changes by Brain Region and Tauopathy"

| **Distance (nm)** | **Normalized Frequency** |
| --- | --- |
| 2.5 | 0.66 |
| 7.5 | 1 |
| 12.5 | 0.9 |
| 17.5 | 0.92 |
| 22.5 | 0.76 |
| 27.5 | 0.86 |
| 32.5 | 0.92 |
| 37.5 | 0.56 |
| 42.5 | 0.56 |
| 47.5 | 0.38 |
| 52.5 | 0.32 |
| 57.5 | 0.36 |
| 62.5 | 0.38 |
| 67.5 | 0.16 |
| 72.5 | 0.2 |
| 77.5 | 0.24 |
| 82.5 | 0.14 |
| 87.5 | 0.2 |
| 92.5 | 0.1 |
| 97.5 | 0.12 |
| 102.5 | 0.08 |
| 107.5 | 0.02 |
| 112.5 | 0.06 |
| 117.5 | 0.04 |
| 122.5 | 0.02 |
| 127.5 | 0 |
| 132.5 | 0.02 |
| 137.5 | 0 |
| 142.5 | 0 |
| 147.5 | 0 |

**Table S1 |** PSD-95 cluster to AMPAR cluster distance data from Fig. 2k.

| **Distance (nm)** | **Normalized Frequency** |
| --- | --- |
| 10 | 0.37143 |
| 30 | 0.85714 |
| 50 | 0.71429 |
| 70 | 1 |
| 90 | 0.91429 |
| 110 | 0.42857 |
| 130 | 0.22857 |
| 150 | 0 |

**Table S2 |** Homer-1 cluster to AMPAR cluster distance data from Fig. 2l.

| **Distance (nm)** | **Paired Correlation (Experimental)** | **Paired Correlation (Randomized)** |
| --- | --- | --- |
| 20 | 1.50021 | 1.15465 |
| 40 | 1.34117 | 1.11987 |
| 60 | 1.18313 | 1.07999 |
| 80 | 1.0549 | 1.03823 |
| 100 | 0.96031 | 1.00369 |
| 120 | 0.90885 | 0.97908 |
| 140 | 0.88924 | 0.97781 |
| 160 | 0.90267 | 0.99851 |
| 180 | 0.94361 | 1.02655 |
| 200 | 1.00097 | 1.05919 |

**Table S3 |** Paired correlation between AMPAR and PSD-95 localizations within a cluster pair from Fig. 2m .

| **Nanodomain Size** | **Frequency Count** |
| --- | --- |
| 5 | 0 |
| 15 | 0 |
| 25 | 0 |
| 35 | 9 |
| 45 | 38 |
| 55 | 77 |
| 65 | 59 |
| 75 | 53 |
| 85 | 46 |
| 95 | 39 |
| 105 | 25 |
| 115 | 11 |
| 125 | 10 |
| 135 | 3 |
| 145 | 3 |
| 155 | 5 |
| 165 | 3 |
| 175 | 2 |
| 185 | 0 |
| 195 | 0 |

**Table S4 |** AMPAR nanodomain size distribution from Fig. 2n.

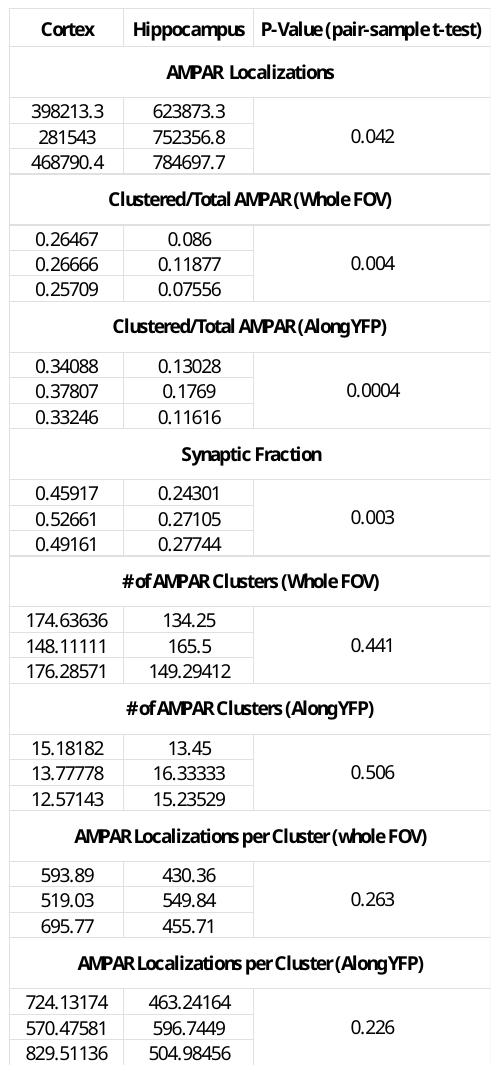

**Table S5 |** Cortex vs Hippocampus comparison datapoints from Fig. 3f-h and Fig. 3k-o . Each datapoint represents average value per mouse.

| **Cortex** | **Hippocampus** |
| --- | --- |
| 0.52629 | 0.12405 |
| 0.41219 | 0.12643 |
| 0.41525 | 0.16657 |
|  | 0.04267 |

**Table S6 |** Clustered/Total AMPARs over whole FOV for intracranially injected CAM2 in Fig. S6b.

| **Cortex** | **Hippocampus** |
| --- | --- |
| 0.443 | 0.23854 |
| 0.29919 | 0.10953 |
| 0.32702 | 0.06098 |
| 0.36914 | 0.16823 |
|  | 0.13904 |
|  | 0.1656 |
|  | 0.1442 |

**Table S7 |** Clustered/Total AMPARs over whole FOV for 5 µM CAM2 labeling in Fig. S6c.

| **Cortical Cultures** | **Hippocampal Cultures** |
| --- | --- |
| 0.06858 | 0.0377 |
| 0.06799 | 0.1765 |
| 0.08194 | 0.13077 |
| 0.13279 | 0.15038 |
| 0.05802 | 0.17316 |
|  | 0.14585 |
|  | 0.19312 |

**Table S8 |** Clustered/Total AMPARs over whole FOV for CAM2 labeling in dissociated neurons in Fig. S6d.

| **WT Cortex** | **WT Hippocampus** | **PS-19 Hippocampus** |
| --- | --- | --- |
| 0.39178 | 0.19346 | 0.27961 |
| 0.46275 | 0.22526 | 0.29002 |
| 0.4562 | 0.2293 | 0.279 |

**Table S9 |** Synaptic fraction of AMPARs as defined within 100 nm of nearest PSD-95 cluster from Fig. S7c, S7d.

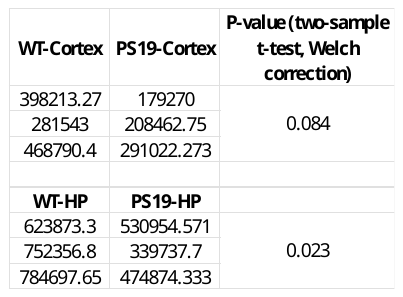

**Table S10 |** AMPAR localizations per FOV comparison datapoints between Wild-type and PS19 mice from Fig. 4c.

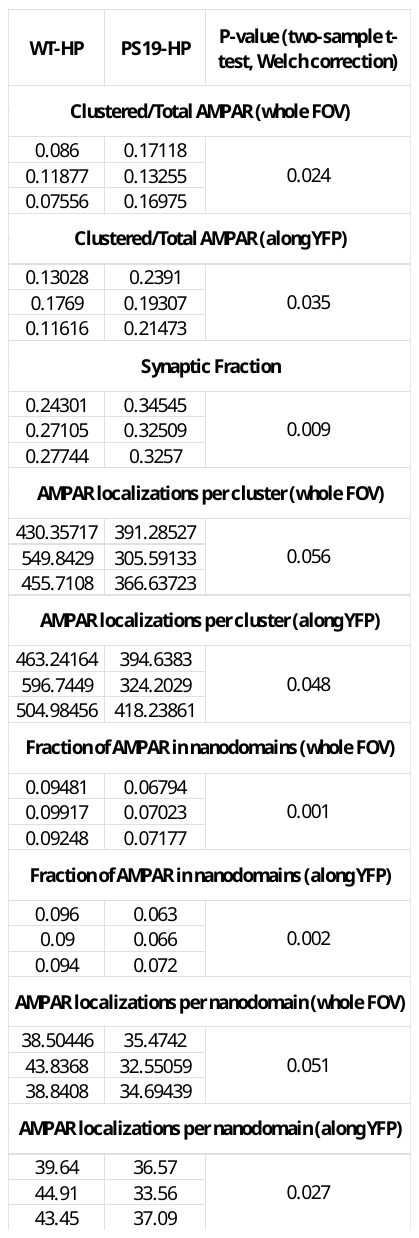

**Table S11 |** Wild-type vs PS19 comparison datapoints from Fig. 4d-l. Each datapoint represents average value per mouse.

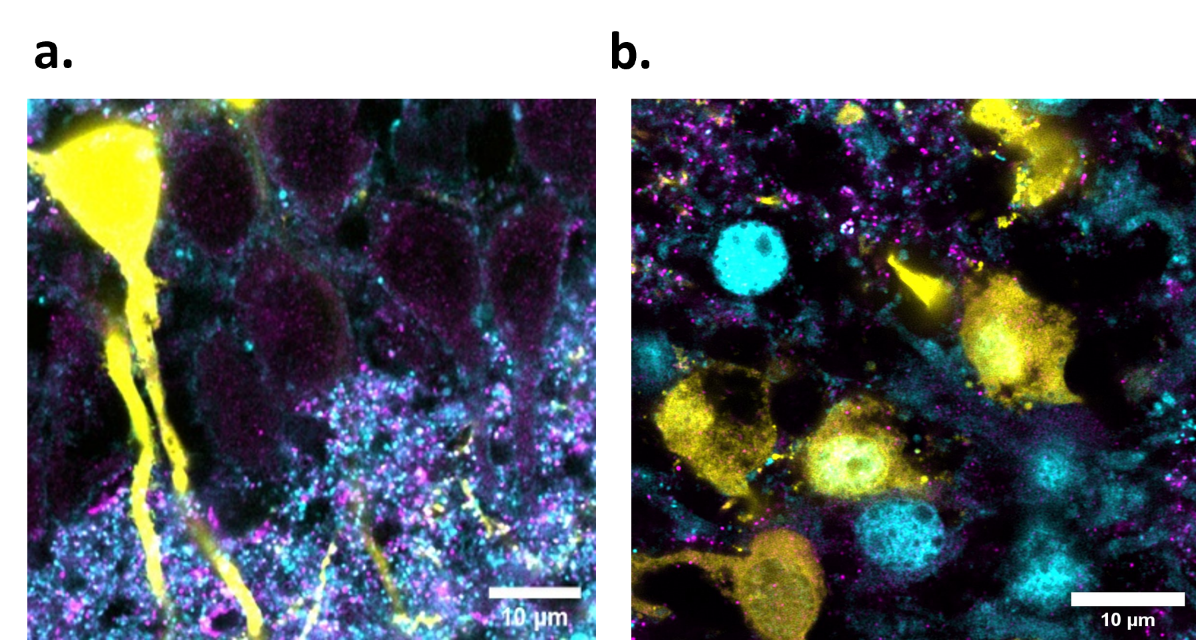

**Figure S1 | Ensuring surface AMPAR labeling.** **a.** Airyscan confocal image of the soma layer in the CA1 region of the hippocampus in a “healthy” brain slice where constant bubbling of 95% O_2_ / 5% CO_2_ was present during CAM2 labeling step. A subset of the neurons is labeled with Thy1-YFP (yellow), along with CAM2-AF 647 (cyan). Homer-1 is also immunostained with CF568 (magenta). No CAM2 labeling is observed in the soma. **b.** Airyscan confocal image of the soma layer in the CA1 region of the hippocampus in a brain slice where 95% O_2_ / 5% CO_2_ was not bubbled during CAM2 labeling. Many soma have CAM2-AF647 (cyan) accumulation in them. Scale bars: 10 µm.

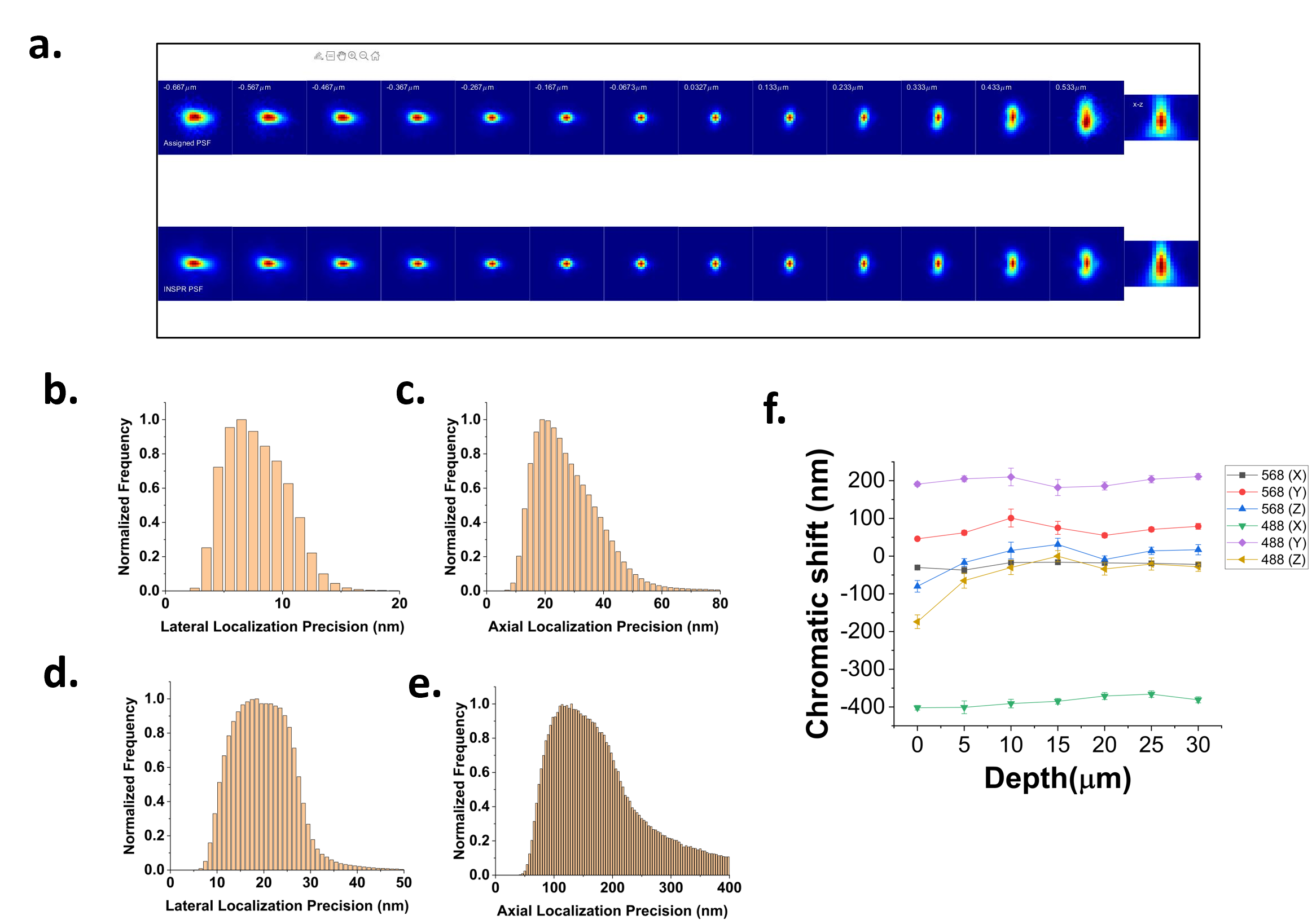

**Figure S2 | Localization Precision and Drift Correction .** **a.** Example of assigned PSF (top row) and the INSPR model PSF (bottom row). Both show good agreement. **b.** Lateral localization precision after INSPR analysis for the image in Fig. 2b. Peak value is ~7 nm. **c.** Axial localization precision after INSPR analysis for the image in Fig. 2b. Peak value is ~22 nm. **d.** Lateral localization precision after conventional elliptical Gaussian fitting analysis using ThunderSTORM plugin in imageJ for the image in Fig. 2b. Peak value is ~17 nm. **e.** Axial localization precision after conventional elliptical Gaussian fitting analysis using ThunderSTORM plugin in imageJ for the image in Fig. 2b. Peak value is ~120 nm. **f.** Chromatic aberration shift for CF568 (denoted by 568) and YFP channels (denoted by 488) in X, Y and Z direction as measured with reference to Alexa Fluor 647 channel.

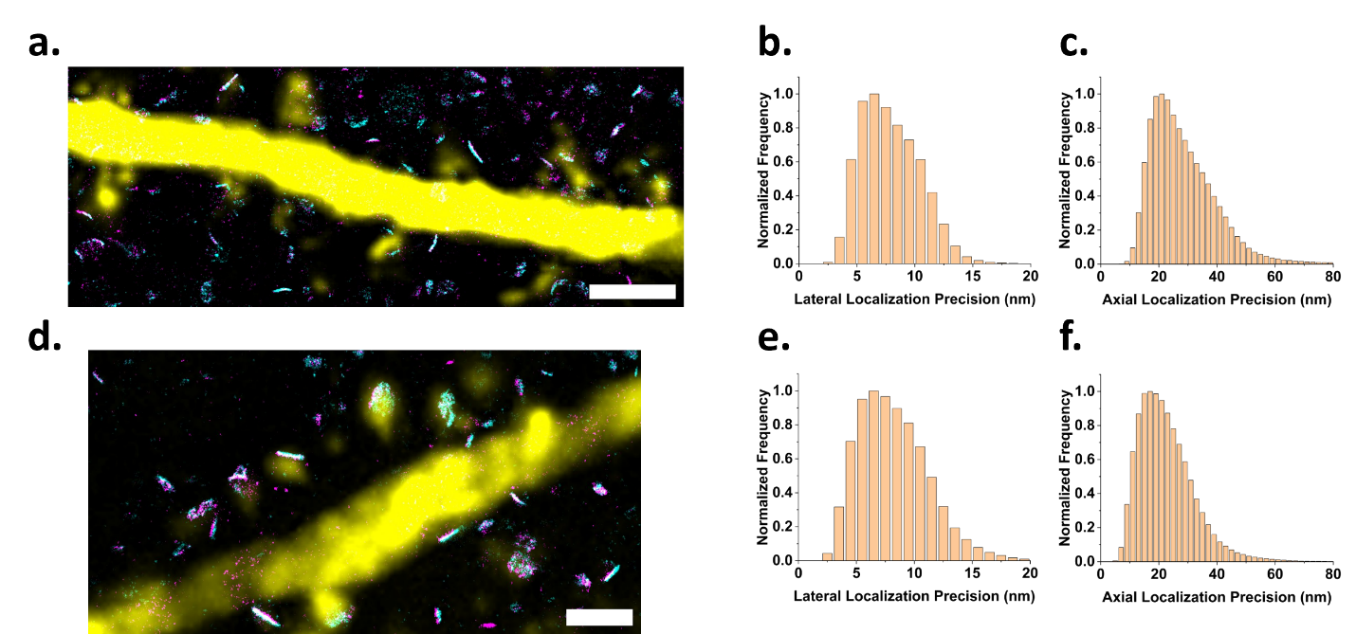

**Figure S3 | Localization precision as a function of depth.** **a.** Example of a dSTORM reconstructed image of CAM2-AF647 and PSD95-CF568 overlayed with Thy1-YFP at an imaging depth of 4 µm. **b.** Lateral localization precision after INSPR analysis of the image in “a”. The peak value is ~7 nm. **c.** Axial localization precision after INSPR analysis of the image in “a”. The peak value is ~22 nm. **d.** Example of dSTORM reconstructed image of CAM2-AF647 and PSD95-CF568 overlayed with Thy1-YFP at an imaging depth of 23 µm. **e.** Lateral localization precision after INSPR analysis of the image in “d”. The peak value is ~8 nm. **f.** Axial localization precision after INSPR analysis of the image in “d”. The peak value is ~25 nm.

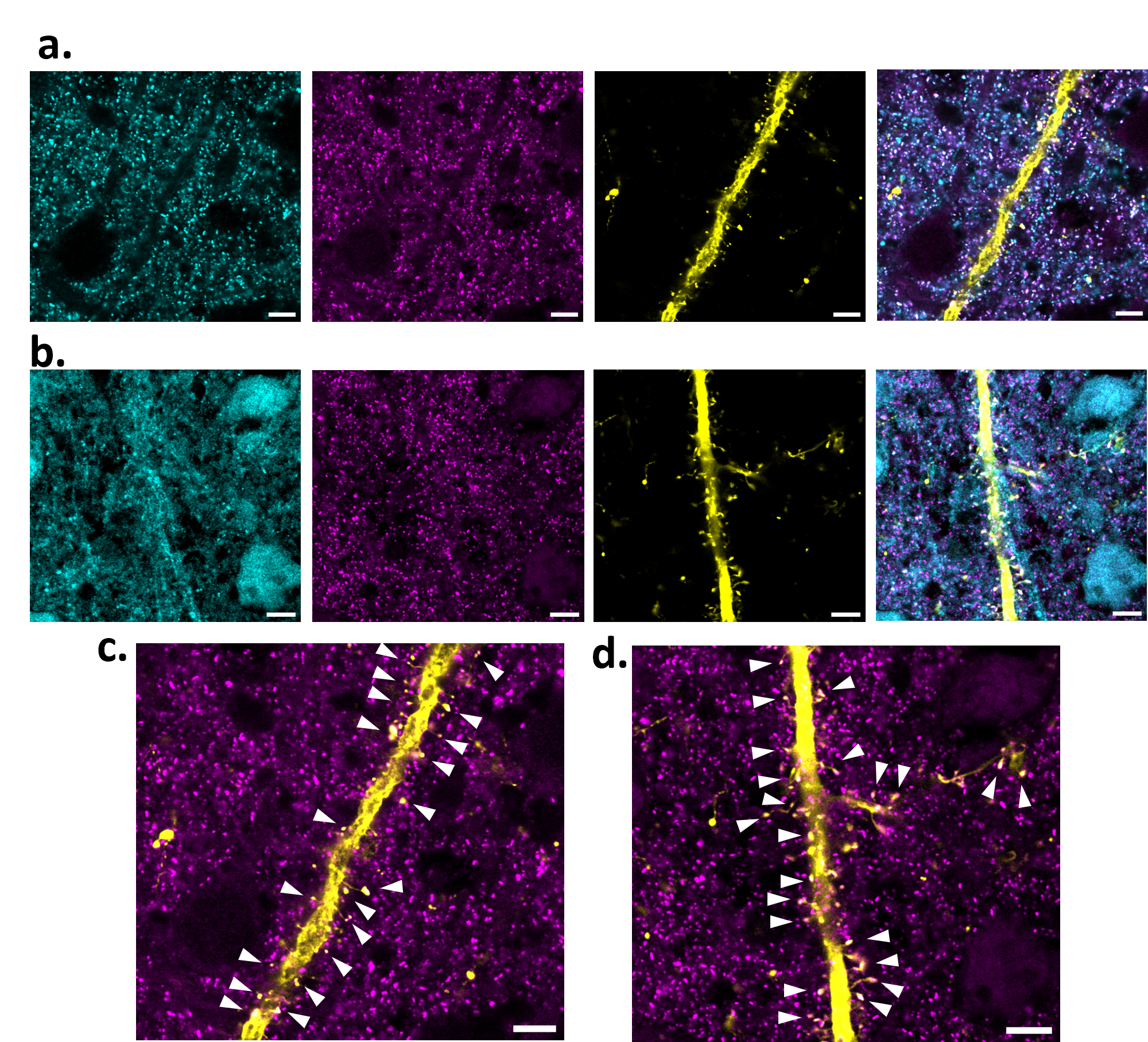

**Figure S4 | PSD-95 labeling comparison in cortex.** **a.** Airyscan confocal images of CAM2-labeled surface AMPAR (cyan), PSD-95 (magenta), Thy1-YFP neuron (yellow) and their composite. **b.** Airyscan confocal images of anti-GluA2/3 antibody-labeled AMPAR (cyan), PSD-95 (magenta), Thy1-YFP neuron (yellow) and their composite. **c.** Composite of PSD-95 (magenta) and Thy1-YFP neuron (yellow) from a. White arrows indicate PSD-95 clusters colocalized with spines. **c.** Composite of PSD-95 (magenta) and Thy1-YFP neuron (yellow) from b. White arrows indicate PSD-95 clusters colocalized with spines.

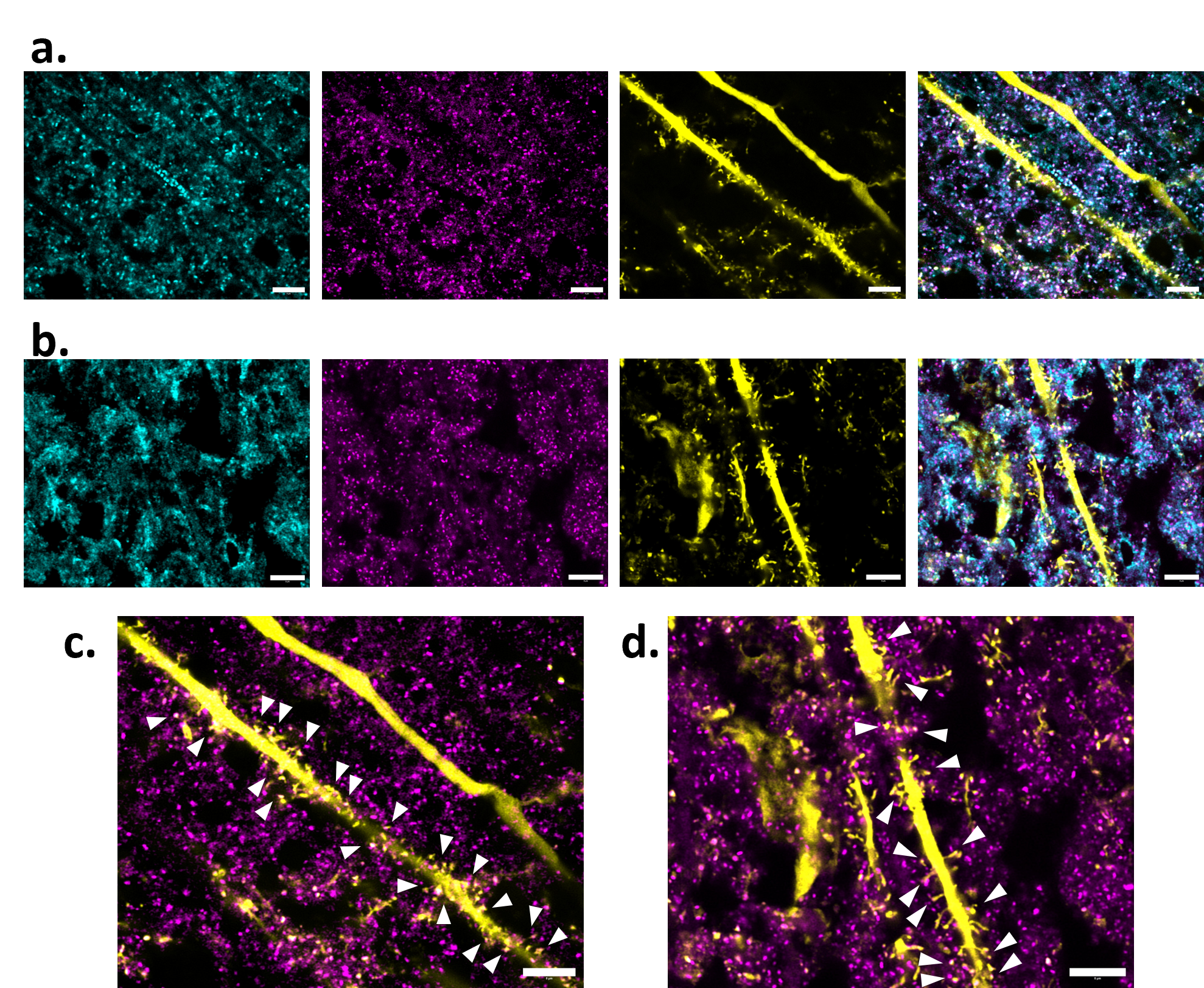

**Figure S5 | PSD-95 labeling comparison in hippocampus.** **a.** Airyscan confocal images of CAM2-labeled surface AMPAR (cyan), PSD-95 (magenta), Thy1-YFP neuron (yellow) and their composite. **b.** Airyscan confocal images of anti-GluA2/3 antibody-labeled AMPAR (cyan), PSD-95 (magenta), Thy1-YFP neuron (yellow) and their composite. **c.** Composite of PSD-95 (magenta) and Thy1-YFP neuron (yellow) from a. White arrows indicate PSD-95 clusters colocalized with spines. **c.** Composite of PSD-95 (magenta) and Thy1-YFP neuron (yellow) from b. White arrows indicate PSD-95 clusters colocalized with spines.

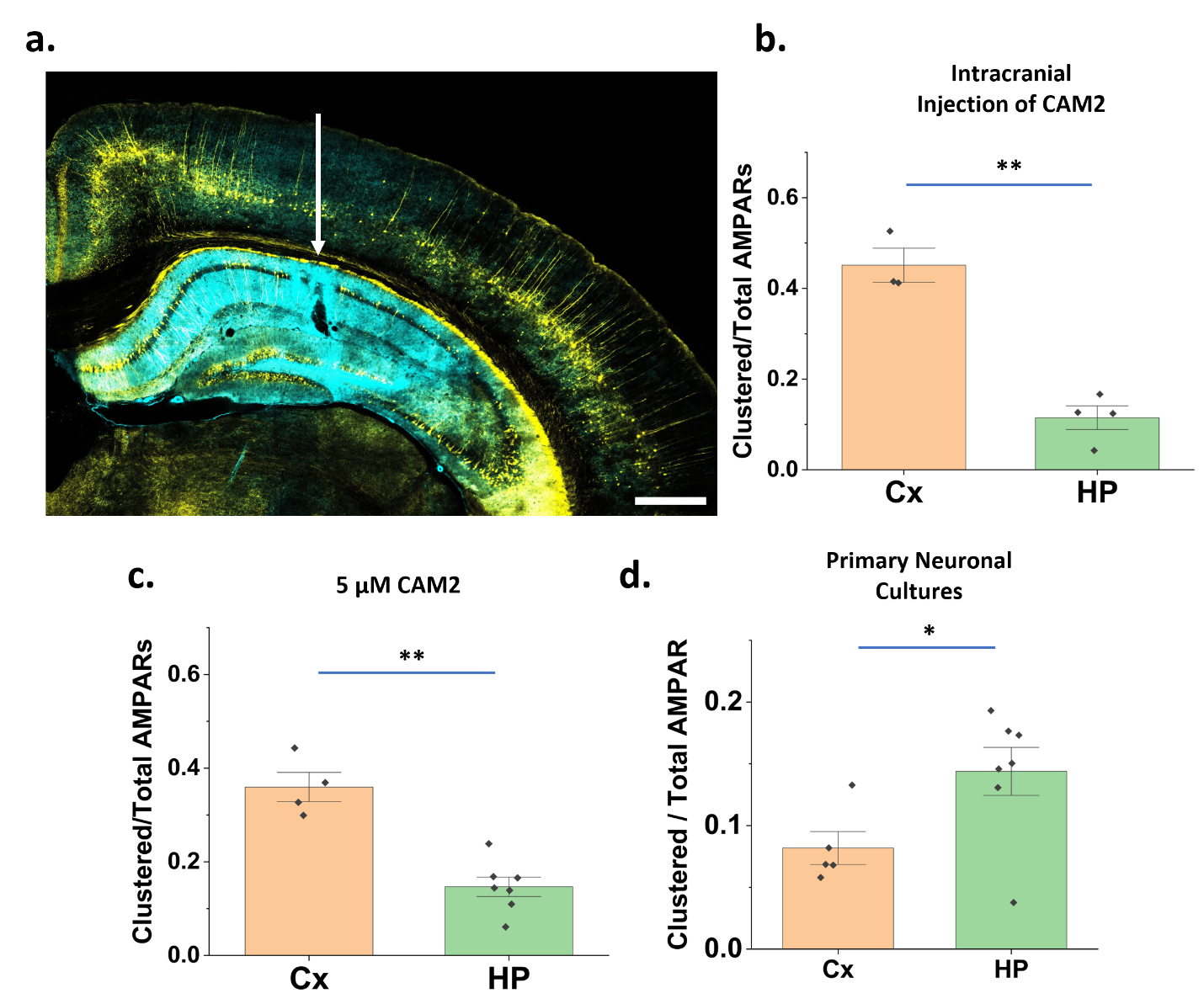

**Figure S6 | CAM2 labeling at higher concentration and dissociated neurons.** **a.** Confocal image of a brain slice after intracranial injection of CAM2-AF647 (50 µM, 2µl) in the CA1 region of the hippocampus of a Thy1-YFP mouse. White arrow indicates the injection site. Strong CAM2 staining (cyan) is observed in the hippocampus. CAM2 also diffuses to stain surface AMPARs in the cortex (although it is dimmer due to lower AMPAR density in the cortex). YFP labeled neurons are shown in yellow. **b.** Fraction of AMPARs found in synaptic clusters across the whole imaging FOV after intracranial injection of CAM2 (n=3-4 FOVs per brain region, p=0.002). **c.** Fraction of AMPARs found in synaptic clusters across the whole imaging FOV at a higher (5 µM) CAM2-AF647 bath labeling concentration (n=4-6 FOVs per brain region, p=0.002). **d.** Fraction of AMPARs found in synaptic clusters across the whole imaging FOV in primary dissociated neuronal cultures from cortex and hippocampus (n=5-7 neurons per brain region from two different cultures, p=0.025).
All error bars depict SE.
b-d: Each datapoint represents one imaging FOV.
**: p<0.01, as determined using two-sample t-test (Welch Correction);
*:p<0.05, as determined using two-sample t-test (Welch Correction);

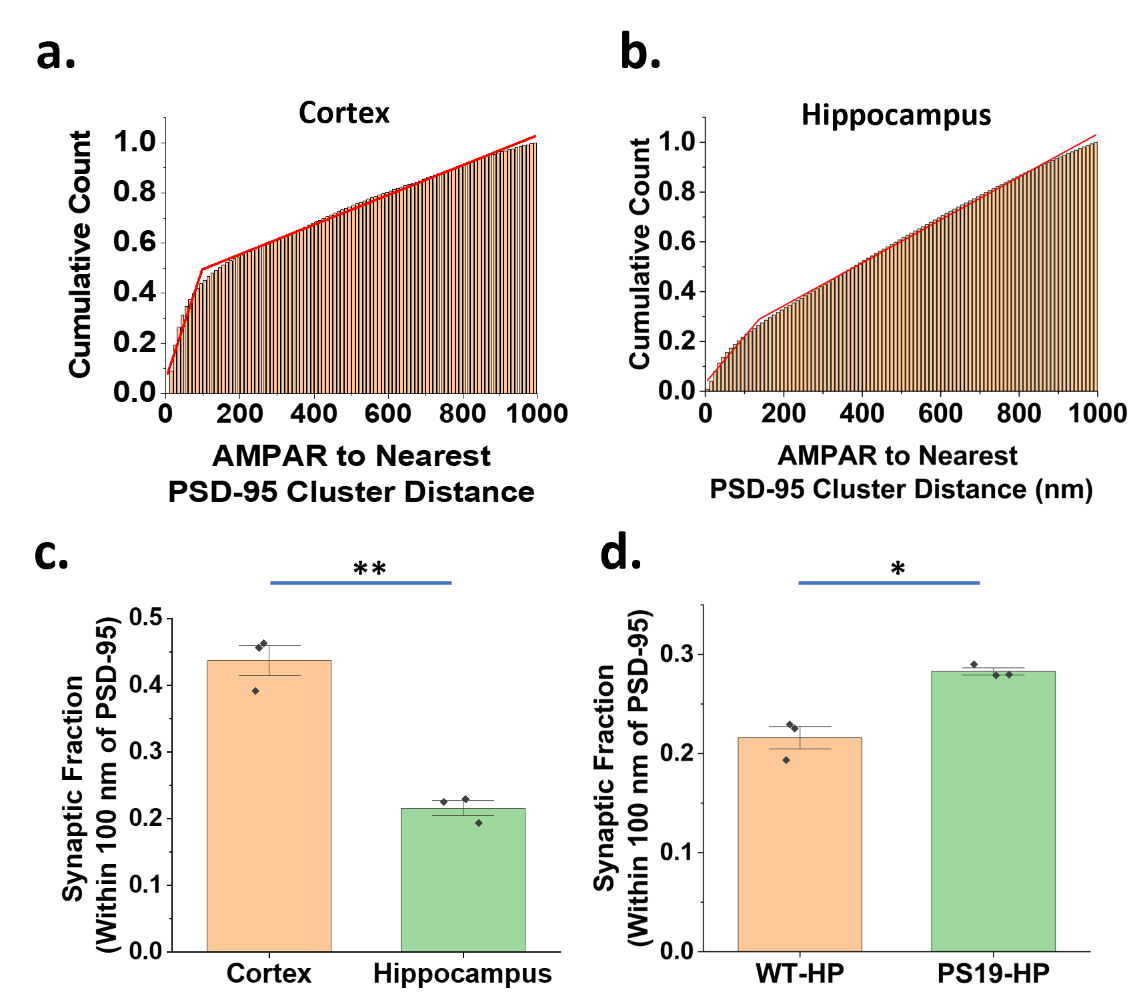

**Figure S7 | Distance threshold for determining synaptic AMPARs.** a. Cumulative frequency distribution of the distance between each AMPAR localization and its nearest PSD-95 cluster in the cortex. The slope change occurs at 97 nm as identified using a piecewise linear fit (R-square=0.99). b. Cumulative frequency distribution of the distance between each AMPAR localization and its nearest PSD-95 cluster in the hippocampus. The slope change occurs at 137 nm as identified using a piecewise linear fit (R-square=0.99). c. Comparison of the fraction of AMPARs lying within 100 nm of a PSD-95 cluster between WT cortex vs WT hippocampus (p=0.003). d. Comparison of the fraction of AMPARs lying within 100 nm of a PSD-95 cluster between WT hippocampus and PS19 hippocampus (p=0.02).
All error bars depict SE.
For a,b: Averaged over 3 mice, at least 3 FoVs per mouse.
For c,d: n=3 mice, at least 3 FoVs per brain region per mouse. Each point represents average value per mouse.

**: p<0.01, as determined using two-sample t-test (Welch Correction);
*p<0.05, as determined using two-sample t-test (Welch Correction);

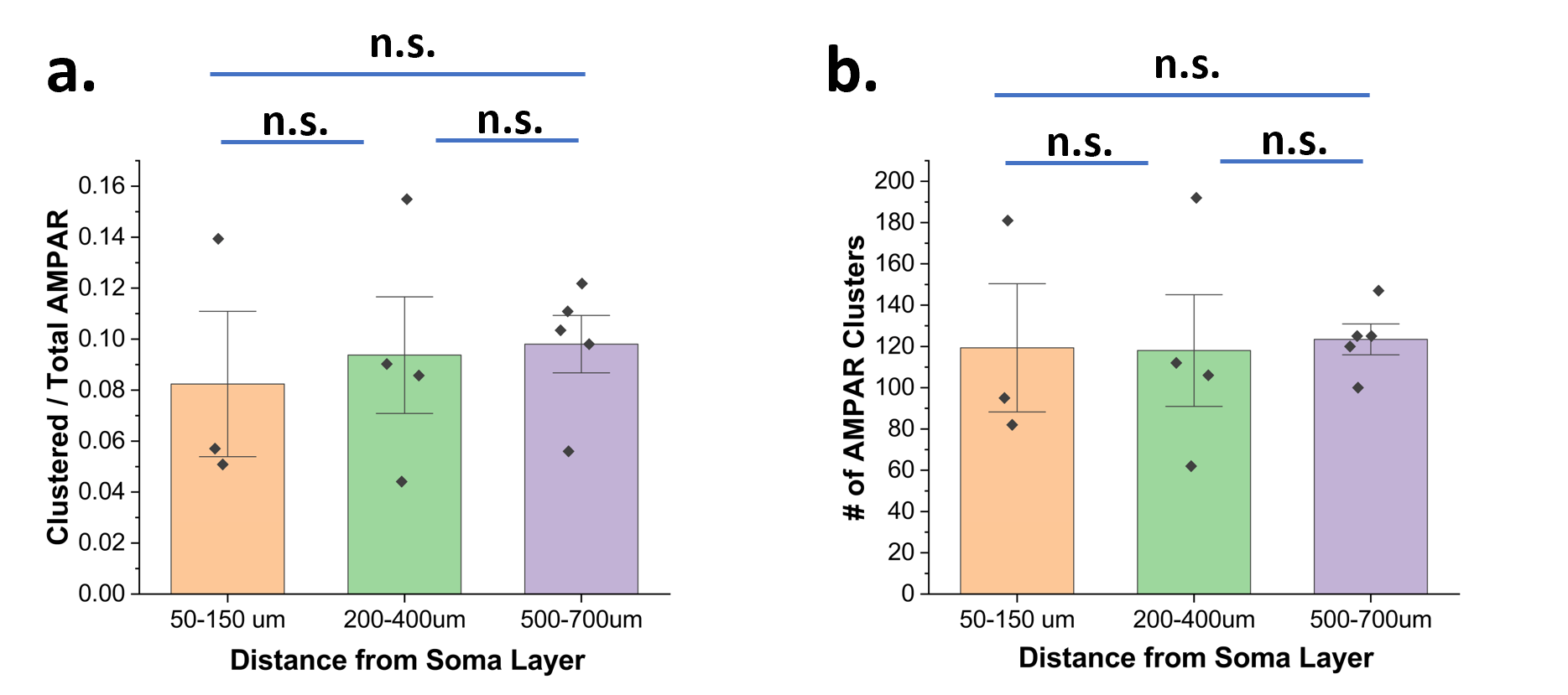

**Figure S8 | AMPAR distribution as a function of distance from CA1 soma layer.** **a.** Fraction of AMPARs found in synaptic clusters across the whole imaging FOV at different distances from the CA1 soma layer. **b.** # of AMPAR clusters per imaging FOV (642 µm^2^) at different distances from the CA1 soma layer.
Each data point represents a separate FOV. Total 3-5 30 µm sections from one Thy1-YFP mouse.
n.s.: p>0.05, as determined using two-sample t-test (Welch Correction).

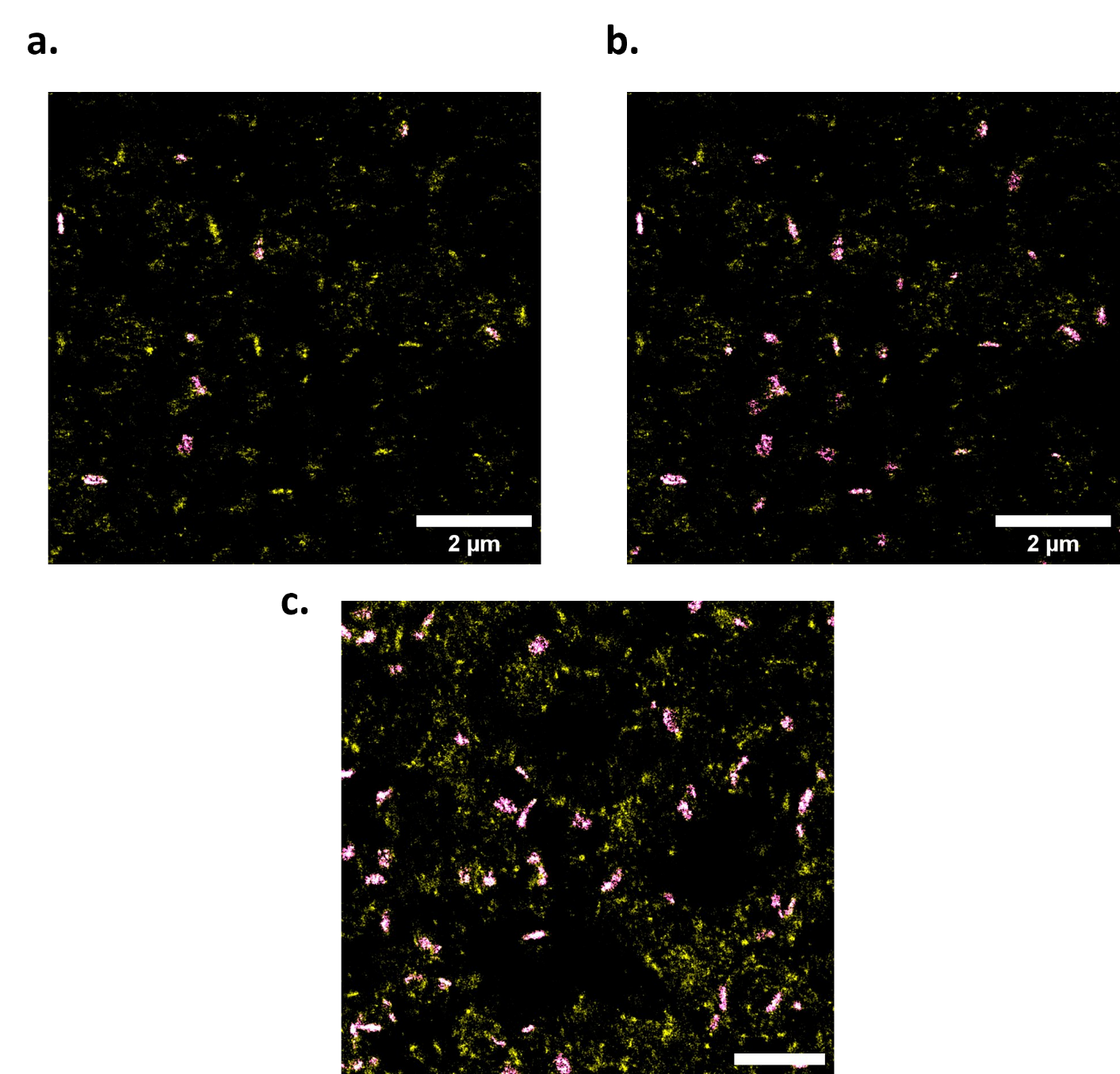

**Figure S9 | Effect of scaling on AMPAR cluster detection in PS19 Hippocampus. a.** With the DBSCAN parameters unchanged, only a few of the high density cluster-like regions in the reconstructed AMPAR localizations (yellow) are detected (magenta). b. After scaling DBSCAN parameters based on average localizations for PS19 mice, most of the clusters are detected (magenta) in the reconstructed data (yellow). c. Cluster detection in a WT hippocampus image for comparison. Without any parameter scaling, clusters are detected very well.

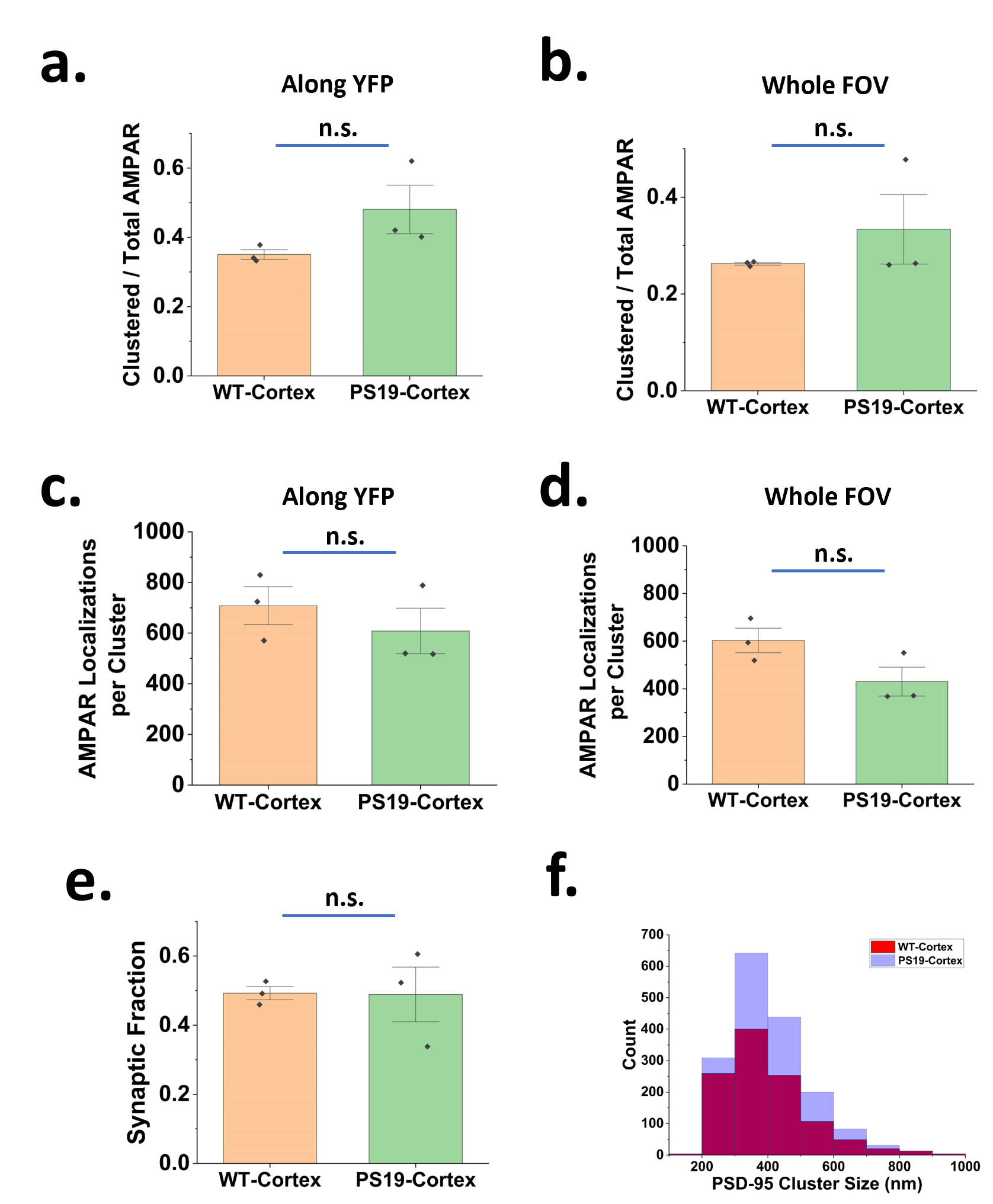

**Figure S10 | WT-cortex vs PS-19-cortex. a.** Fraction of AMPARs on a YFP neuron found in clusters. **b,** Fraction of AMPARs in the whole FoV found in clusters. **c,** AMPAR localizations per cluster along a YFP neuron **d,** AMPAR localizations per cluster over whole FOV. **e,** Fraction of AMPARs lying within 140 nm of a PSD-95 cluster. f, Histograms of PSD-95 cluster size distribution for WT cortex (red) and PS19 cortex (blue).
All error bars depict SE.
For a-d and f: n=3 mice each for WT and PS19, at least 6 FoVs per mouse per brain region. Each point represents average value per mouse.
For e: n=3 mice each for WT and PS19, at least 3 FoVs per mouse. Each point represents average value per mouse.
**: p<0.01, as determined using two-sample t-test (Welch Correction);
*p<0.05, as determined using two-sample t-test (Welch Correction);
n.s.: p>0.05 as determined using two-sample t-test (Welch Correction).
