## Supplementary Movies for "Synaptic Organization of Surface AMPARs Changes by Brain Region and Tauopathy"

### Slide 1
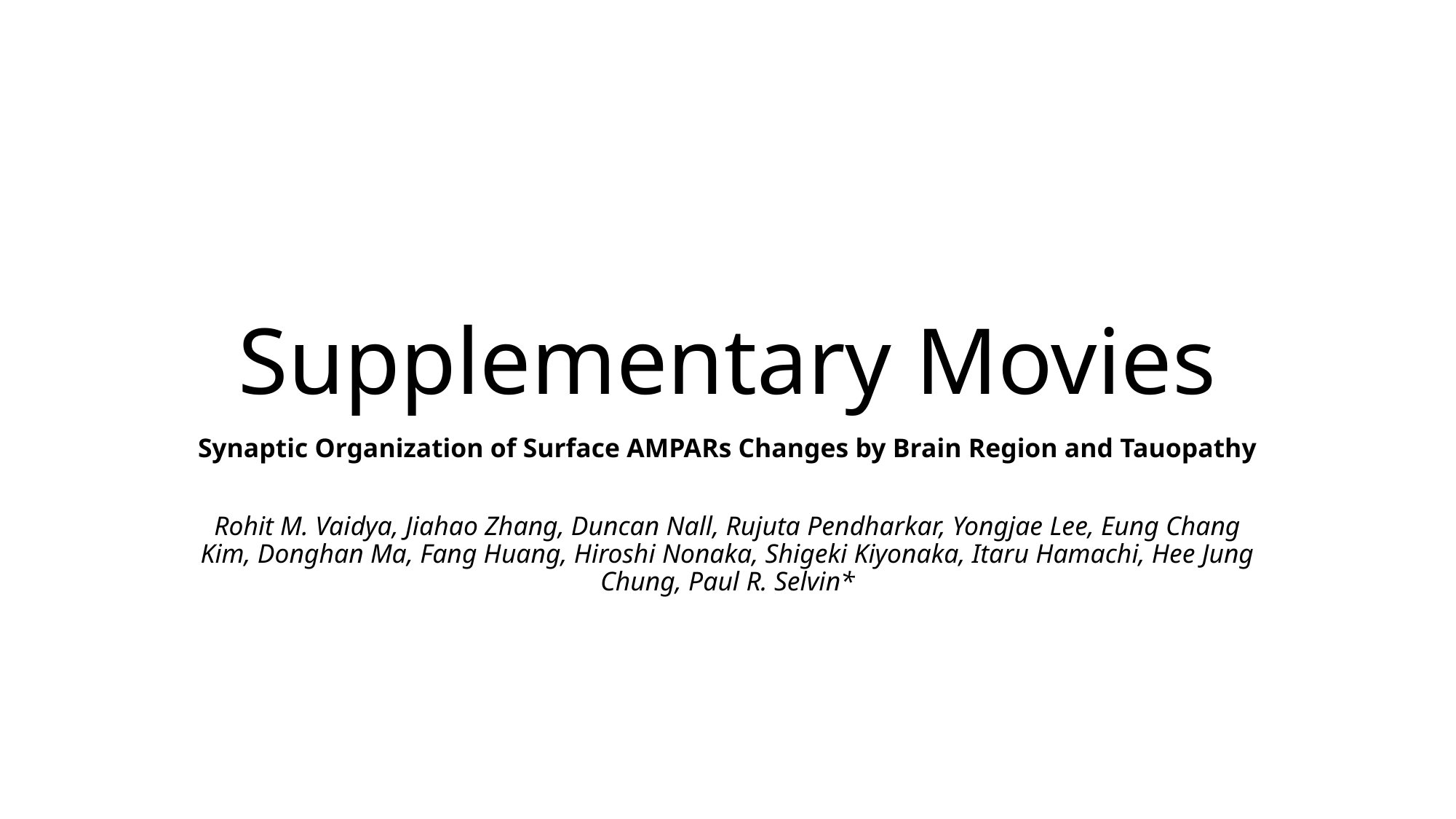

#### Supplementary Movies
Synaptic Organization of Surface AMPARs Changes by Brain Region and Tauopathy
Rohit M. Vaidya, Jiahao Zhang, Duncan Nall, Rujuta Pendharkar, Yongjae Lee, Eung Chang Kim, Donghan Ma, Fang Huang, Hiroshi Nonaka, Shigeki Kiyonaka, Itaru Hamachi, Hee Jung Chung, Paul R. Selvin*

### Slide 2
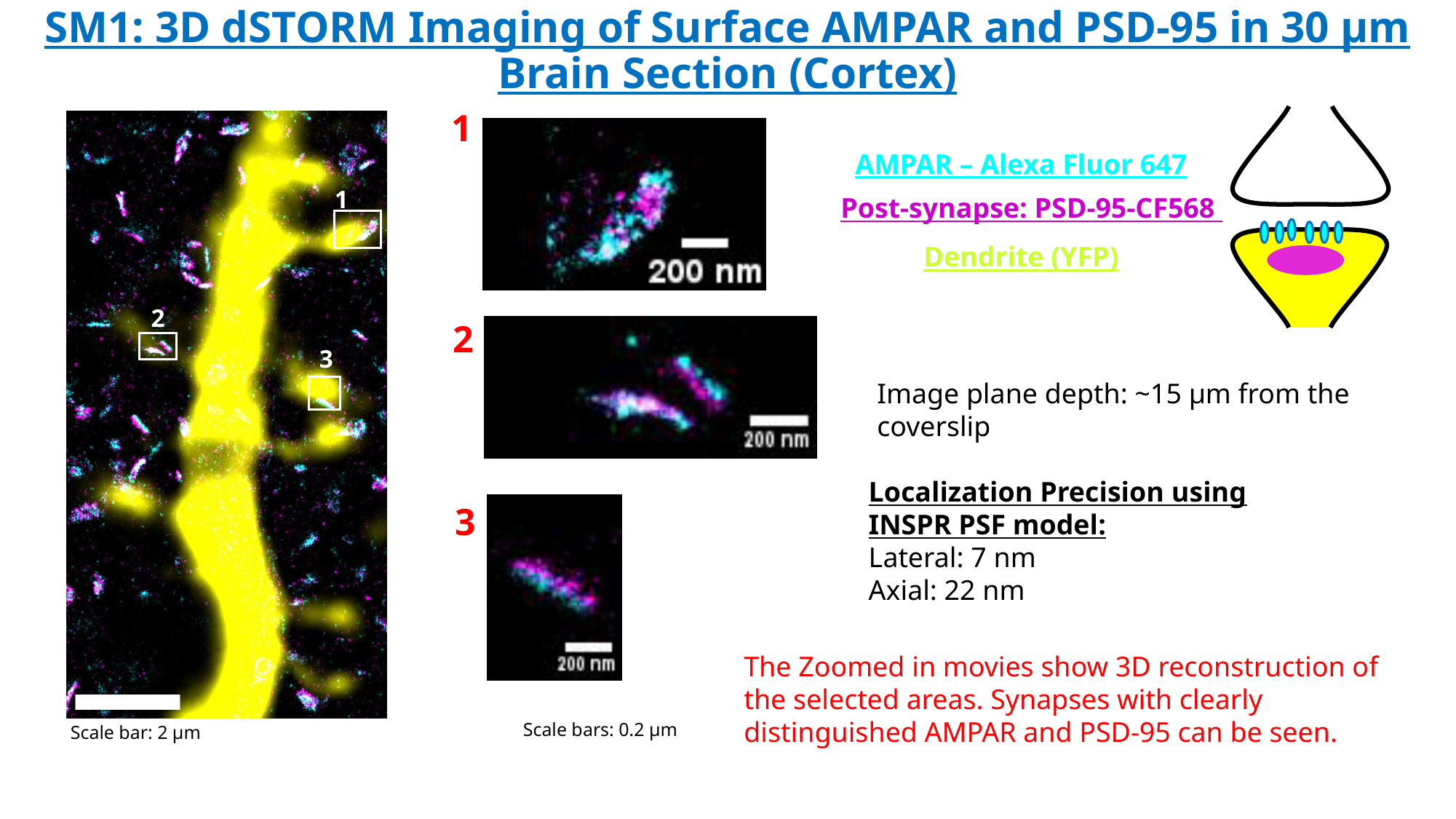

#### SM1: 3D dSTORM Imaging of Surface AMPAR and PSD-95 in 30 µm Brain Section (Cortex)
1
AMPAR – Alexa Fluor 647
Post-synapse: PSD-95-CF568
Dendrite (YFP)
1
2
2
3
Image plane depth: ~15 µm from the coverslip
Localization Precision using INSPR PSF model:
Lateral: 7 nm
Axial: 22 nm
3
The Zoomed in movies show 3D reconstruction of the selected areas. Synapses with clearly distinguished AMPAR and PSD-95 can be seen.
Scale bars: 0.2 µm
Scale bar: 2 µm

### Slide 3
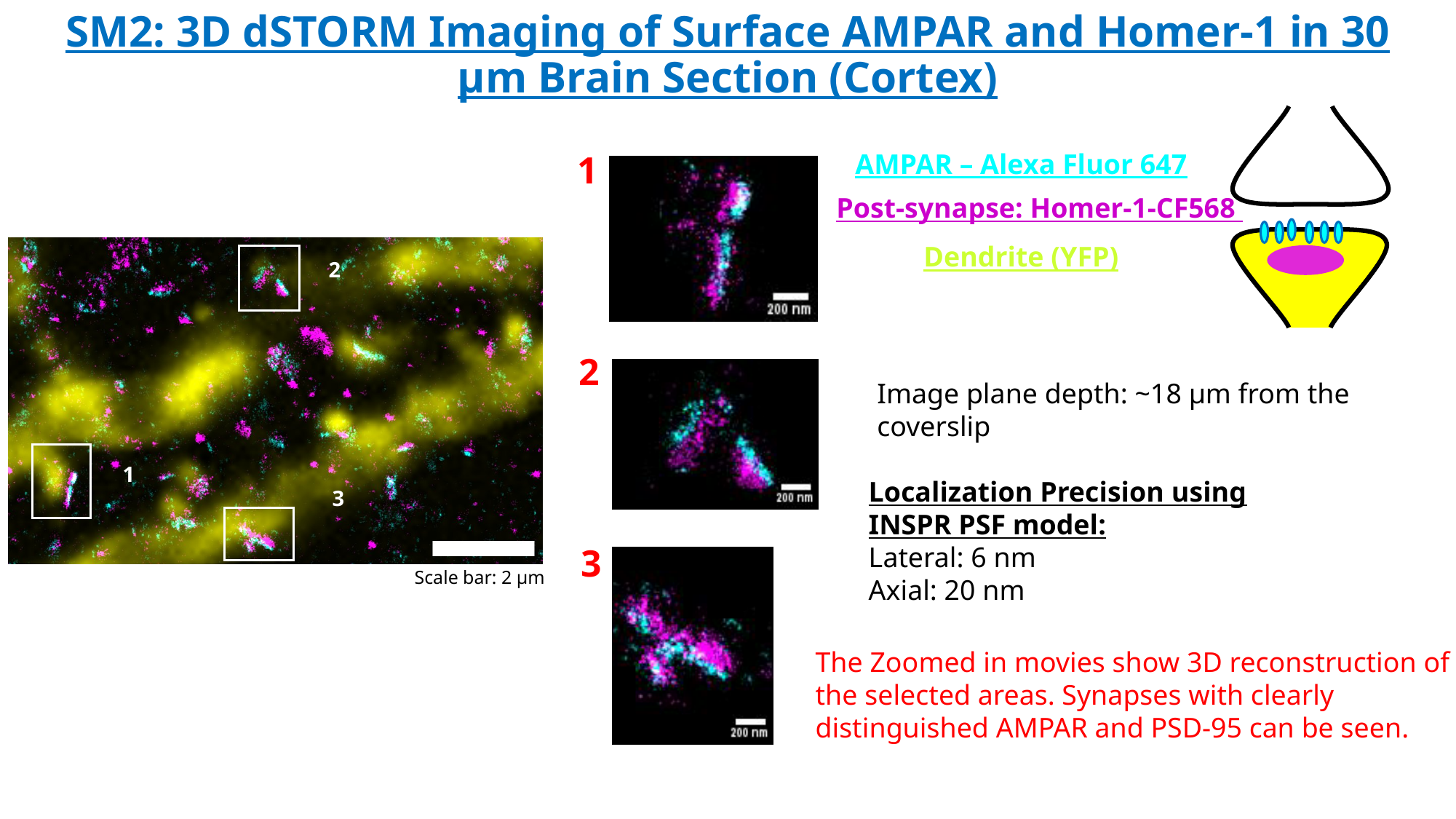

#### SM2: 3D dSTORM Imaging of Surface AMPAR and Homer-1 in 30 µm Brain Section (Cortex)
AMPAR – Alexa Fluor 647
Post-synapse: Homer-1-CF568
Dendrite (YFP)
1
2
1
3
Scale bar: 2 µm
2
Image plane depth: ~18 µm from the coverslip
Localization Precision using INSPR PSF model:
Lateral: 6 nm
Axial: 20 nm
3
The Zoomed in movies show 3D reconstruction of the selected areas. Synapses with clearly distinguished AMPAR and PSD-95 can be seen.
